# Mriyaviruses: Small Relatives of Giant Viruses

**DOI:** 10.1101/2024.02.29.582850

**Authors:** Natalya Yutin, Pascal Mutz, Mart Krupovic, Eugene V. Koonin

## Abstract

The phylum *Nucleocytoviricota* consists of large and giant viruses that range in genome size from about 100 kilobases (kb) to more than 2.5 megabases. Here, using metagenome mining followed by extensive phylogenomic analysis and protein structure comparison, we delineate a distinct group of viruses with double-stranded (ds) DNA genomes in the range of 35-45 kb that appear to be related to the *Nucleocytoviricota.* In phylogenetic trees of the conserved double jelly-roll major capsid proteins (MCP) and DNA packaging ATPases, these viruses do not show affinity to any particular branch of the *Nucleocytoviricota* and accordingly would comprise a class which we propose to name “*Mriyaviricetes*” (after Ukrainian Mriya, dream). Structural comparison of the MCP suggests that, among the extant virus lineages, mriyaviruses are the closest one to the ancestor of the *Nucleocytoviricota*. In the phylogenetic trees, mriyaviruses split into two well-separated branches, the family *Yaraviridae* and proposed new family “*Gamadviridae*”. The previously characterized members of these families, Yaravirus and Pleurochrysis sp. endemic viruses, infect amoeba and haptophytes, respectively. The genomes of the rest of the mriyaviruses were assembled from metagenomes from diverse environments, suggesting that mriyaviruses infect various unicellular eukaryotes. Mriyaviruses lack DNA polymerase, which is encoded by all other members of the *Nucleocytoviricota,* and RNA polymerase subunits encoded by all cytoplasmic viruses among the *Nucleocytoviricota*, suggesting that they replicate in the host cell nuclei. All mriyaviruses encode a HUH superfamily endonuclease that is likely to be essential for the initiation of virus DNA replication via the rolling circle mechanism.

**Importance:** The origin of giant viruses of eukaryotes that belong to the phylum *Nucleocytoviricota* is not thoroughly understood and remains a matter of major interest and debate. Here we combine metagenome database searches with extensive protein sequence and structure analysis to describe a distinct group of viruses with comparatively small genomes of 35-45 kilobases that appears to comprise a distinct class within the phylum *Nucleocytoviricota* that we provisionally named *“Mriyaviricetes”.* Mriyaviruses appear to be the closest identified relatives of the ancestors of the *Nucleocytoviricota.* Analysis of proteins encoded in mriyavirus genomes suggest that they replicate their genome via the rolling circle mechanism that is unusual among viruses with double-stranded DNA genomes and so far not described for members of *Nucleocytoviricota*.

## Introduction

The phylum *Nucleocytoviricota* (informally also known as NCLDV, Nucleo-Cytoplasmic Large DNA Viruses) unites large and giant viruses that range in genome size from about 100 kilobases (kb) to more than 2.5 megabases (1, 2). The origin of the giant viruses has been hotly debated, and scenarios of their reductive evolution from cellular life forms, possibly, a “fourth domain of life”, have been actively discussed (3–6). However, genome evolution reconstruction based on phylogenies of conserved viral genes clearly indicates that the giant viruses (operationally defined as those with genomes larger than 500 kb) within *Nucleocytoviricota* evolved from smaller viruses on multiple, independent occasions, capturing genes from their eukaryotic hosts, bacteria, and other viruses (2, 7–10).

Thus, genomic gigantism appears to be a derived feature among the *Nucleocytoviricota,* with the implication that minimalistic members of this phylum, perhaps, resembling the ancestral forms, potentially could be discovered. Indeed, two groups of viruses with comparatively small genomes apparently belonging to *Nucleocytoviricota* have been recently reported. The first of these consists of viruses infecting crustacea that have been assigned to the putative family “*Mininucleoviridae*”, with the genomes in the range of 70 to 74 kb (11). The protein sequences of mininucleoviruses are highly divergent, but nevertheless, phylogenetic analysis of hallmark genes that are conserved across the *Nucleocytoviricota* confidently places them within the order *Pimascovirales* (11). Thus, the comparatively small genome size in the viruses of this family is a derived character resulting from reductive evolution. The second group includes viruses with even smaller genomes and represented by the family *Yaraviridae*, currently including a single representative, Yaravirus, with the genome of about 45 kb (12, 13), and *Pleurochrysis* sp. endemic viruses (PEV), with genomes of about 35 kb. Along with some *Phaeocystis*-related metagenomic contigs, PEV also have been independently referred to as “NCLDV-like dwarf viruses” (NDDV) (14). Most of the proteins of these viruses have no readily detectable homologs such that even their relationships with the *Nucleocytoviricota* remained uncertain.

We sought to characterize in detail the smallest putative members of the *Nucleocytoviricota* and their relationship with other viruses in this phylum. To this end, we searched genomic and metagenomic databases for homologs of the double jelly-roll (DJR) major capsid proteins (MCP) of Yaravirus and PEV. These searches led to the identification of an expansive group of viruses with genomes in the 35-45 kilobase (kb) range, with two subgroups, one related to Yaravirus and the other one to PEV. We performed a comprehensive phylogenomic analysis of these virus genomes and identified several conserved proteins shared with other members of *Nucleocytoviricota* as well as a set of proteins conserved specifically within this group. Phylogenetic analysis of the conserved proteins supported the monophyly of this group but failed to detect specific affinity with any other group within *Nucleocytoviricota.* We therefore suggest that these viruses should be classified as a class within the phylum *Nucleocytoviricota* which we propose to name “*Mriyaviricetes*” (after the Ukrainian Mriya, dream). Structural comparisons of the MCPs suggest that mriyaviruses could be the extant group of viruses most closely related to the common ancestor of the *Nucleocytoviricota*.

## Results

### Identification of mriyaviruses, a distinct group of viruses with small genomes related to Nucleocytoviricota

We sought to identify members or relatives of the phylum *Nucleocytoviricota* with small genomes and to this end searched the publicly available genomic and metagenomic sequence databases for proteins with significant similarity to the MCPs of “*Mininucleoviridae*”, Yaravirus, and NDDV. No proteins significantly similar to mininucleovirus MCPs were detected, but the searches initiated with the sequences of the MCPs of Yaravirus and NDDV produced about 2,000 significant hits. These protein sequences were clustered, cluster representatives were aligned with MCPs of representatives of the major groups of *Nucleocytoviricota*, and a phylogenetic tree was constructed from the alignment. In this tree, about 200 MCP sequences formed a strongly supported clade that included Yaravirus and NDDV, indicative of the monophyly of these viruses (Figure 1a). The contigs encoding these predicted MCPs originated from various environments, including marine, freshwater, and soil microbiomes (Supplementary Table S1), suggesting a broad host range. We named this virus group “Mriyaviruses” (from Ukrainian ‘mriya’ – dream). The mriyavirus clade split into two distinct branches, one of which included Yaravirus, and the other one included the NDDV. We denoted the former group *Yaraviridae*, after the already approved virus family (13), and the latter group “*Gamadviridae*”, a putative new family (from Hebrew ‘gamad’, dwarf). *Yaraviridae* is a far more diverse group than “*Gamadviridae*” and potentially might be elevated in taxonomic rank and split into several families in the future.

**Figure 1.**
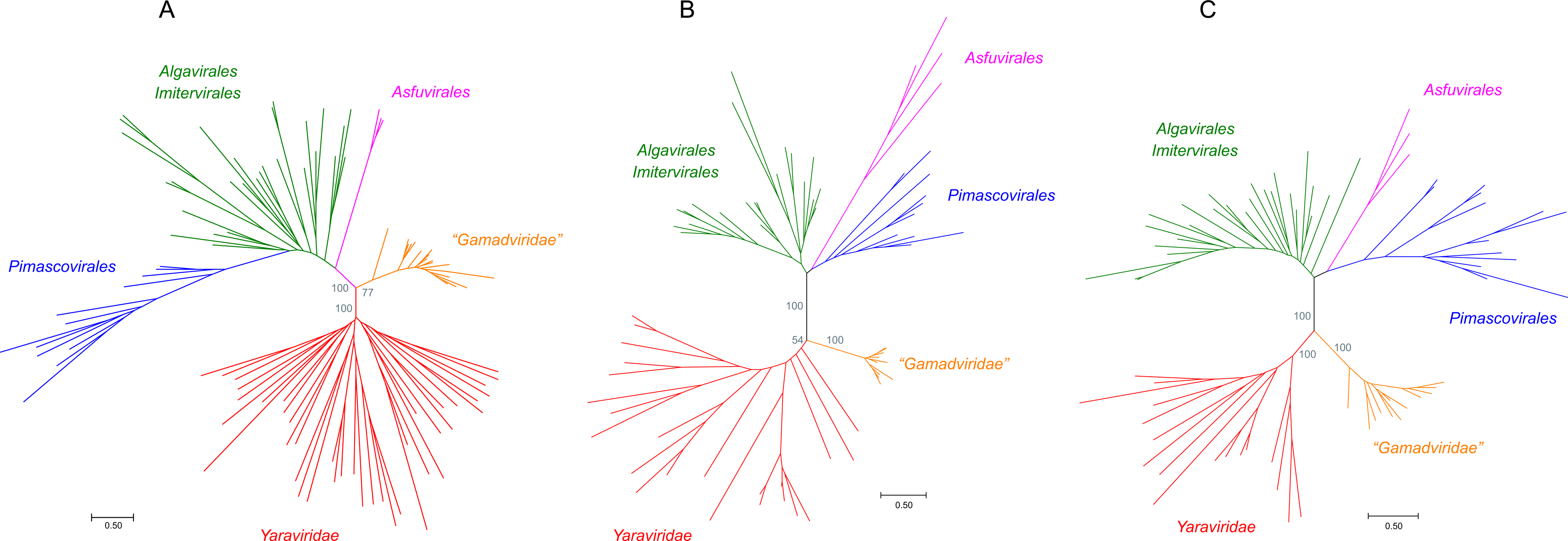
Phylogenetic trees of proteins conserved in mriyaviruses and the rest of the members of *Nucleocytoviricota*. A, Major Capsid Protein (MCP); B, DNA packaging ATPase (ATPase); C, Virus Late Transcription Factor 3 (VLTF3). The IQTree bootstrap values are indicated for the key branches. The trees in newick format are accessible at https://ftp.ncbi.nih.gov/pub/yutinn/mriya_2024.

Using the MCP tree as a guide, 60 representative genomes and long contigs were selected for detailed analysis based on the length and diversity coverage. Among the members of the *Yaraviridae*, there were several long contigs containing direct terminal repeats, suggesting that the respective genomes are complete and furthermore are circular or terminally redundant (genetically circular) (Figure 2). The predicted protein sequences encoded by the 60 representative mriyaviruses genomes were clustered and annotated using HHpred and CDD searches and structures of selected proteins of interest (see below) were modeled using AlphaFold 2 (AF2) or ColabFold.

**Figure 2.**
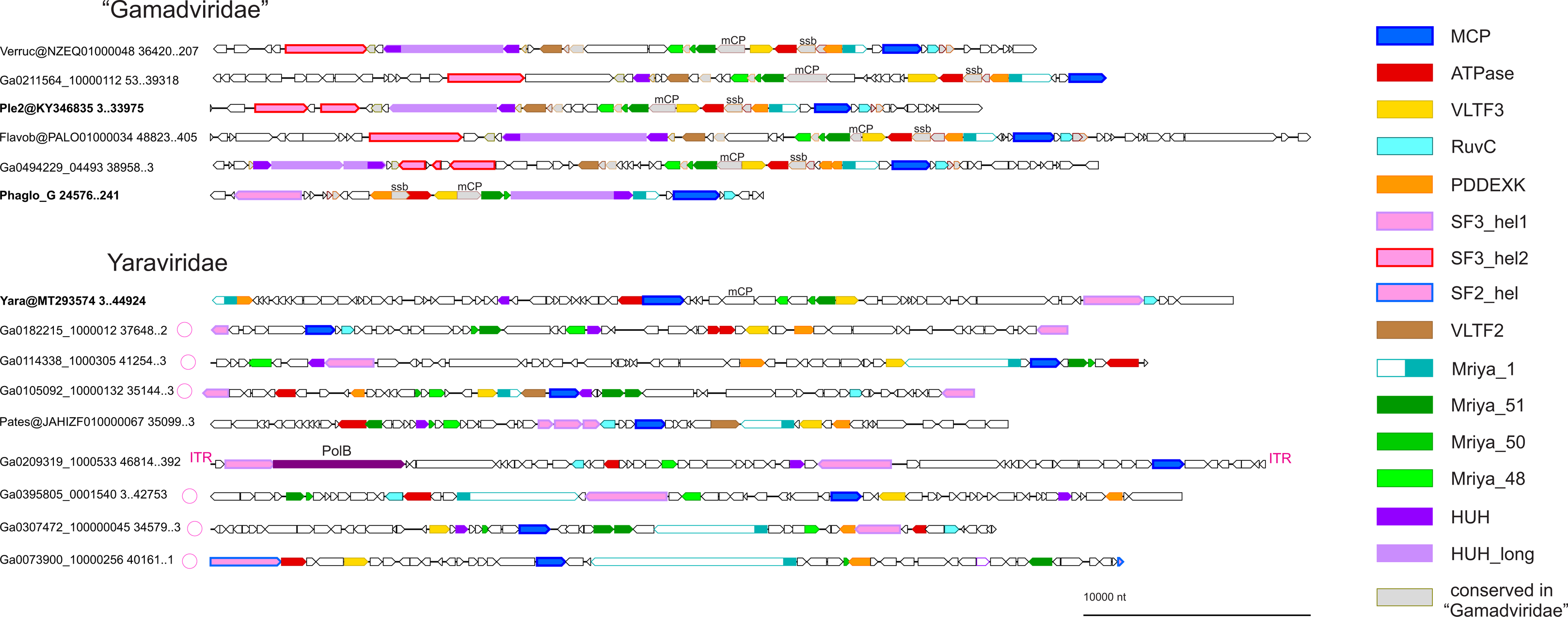
Genome maps of selected mriyaviruses. Genes with predicted functions are shown by color-coded block arrows. Circles near contig names indicate contigs with direct terminal repeats. Abbreviations: ITR, inverted terminal repeats; PolB, family B DNA polymerase; mCP, minor capsid protein; ssb, single strand DNA binding protein; MCP, Major Capsid Protein; ATPase, DNA packaging ATPase; VLTF3, virus late transcription factor 3, RuvC, RuvC-like Holliday junction resolvase homologous to poxvirus A22 resolvase; PDDEXK, PDDEXK superfamily endonuclease; VLTF2, virus late transcription factor 2; Mriya_1, conserved domain homologous to Yaravirus gene 1; Mriya_51, Yaravirus gene 51 homolog; Mriya_50, Yaravirus gene 50 homolog; Mriya_48, Yaravirus gene 48 homolog; HUH, mriyavirus HUH endonuclease; HUH_long, conserved gamadvirus protein containing a C-terminal domain homologous to mriyavirus HUH endonuclease. Genome maps of all 60 mriyavirus representative genomes are available at https://ftp.ncbi.nih.gov/pub/yutinn/mriya_2024.

### Phylogenomics of mriyaviruses

We identified 12 (predicted) proteins that were conserved in (nearly) all mriyaviruses and, in addition, 10 proteins that were conserved in all members of “*Gamadviridae*” but lacked detectable homologs in *Yaraviridae* (Figure 3, Supplementary Table S1, and Table 1). Among the 12 conserved proteins that unite the mriyaviruses, 5 are homologous to proteins that are conserved across the phylum *Nucleocytoviricota*, namely, MCP, DNA packaging ATPase (ATPase), viral late gene transcription factor 2 (VLTF2), viral late gene transcription factor 3 (VLTF3) and the RuvC-like Holliday junction resolvase (RuvC). The conservation of VLTF2 and VLTF3 in itself appears diagnostic of the affinity of mriyaviruses with the *Nucleocytoviricota* because homologs of these proteins were not detectable outside this virus phylum.

**Figure 3.**
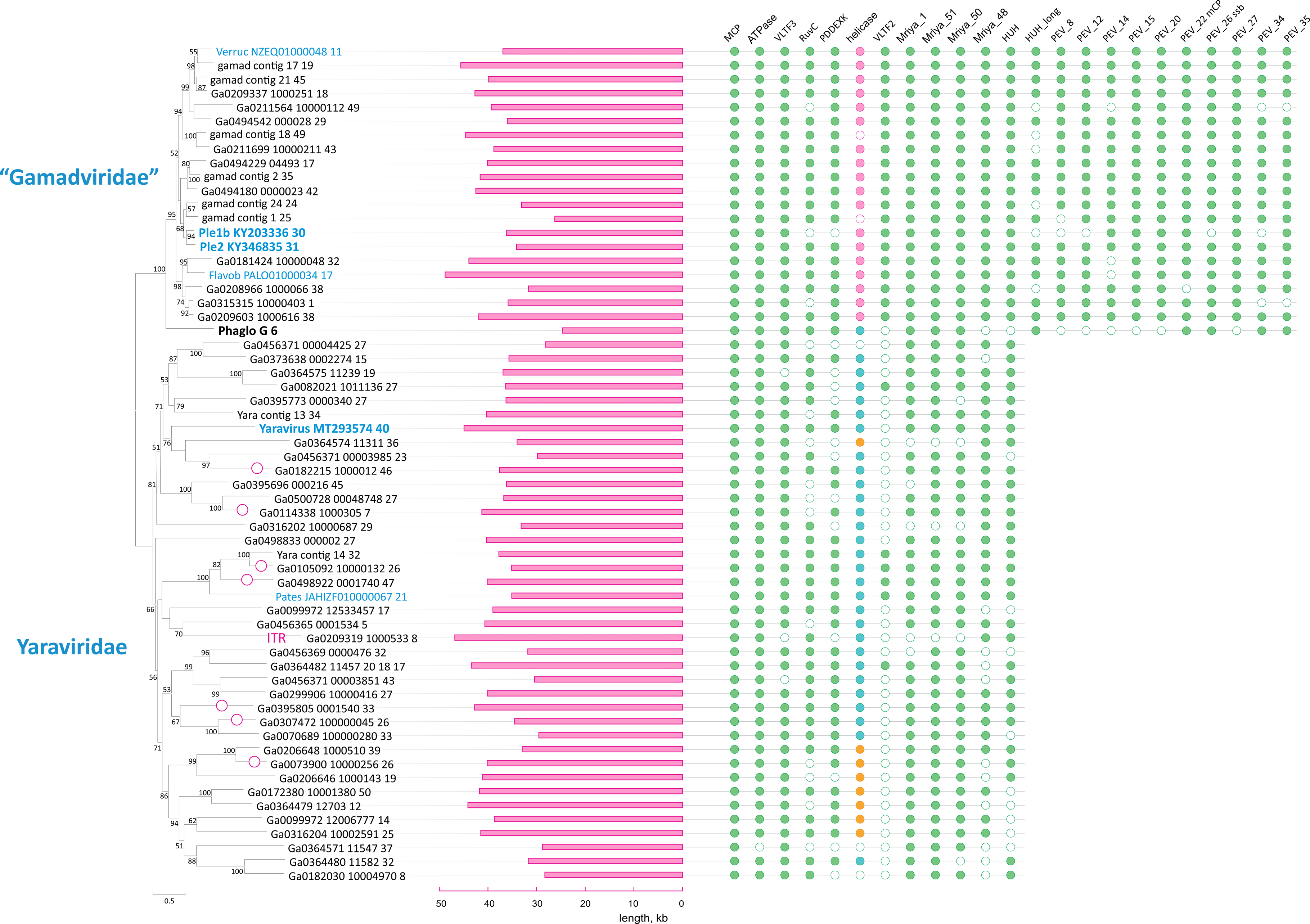
Patterns of protein presence-absence in mriyaviruses. The MCP tree was rooted between *Yaraviridae* and “*Gamadviridae*” for visualization. Circles at branches indicate contigs with terminal repeats. Genomes retrieved from GenBank are denoted with blue font. The middle panel shows genome length. Conserved proteins are abbreviated as in Figure 2. The coloring in the helicase column indicates: turquoise, SF3 family helicase (SF3_hel1 group); pink, SF3 family helicase (SF3_hel2); orange, SF2 family helicase (SF2_hel).

**Table 1.** Proteins conserved in *Mriyaviricetes*.

| conserved protein |  | *Yaravirus brasiliensis* protein ID |  | P. endemic virus 2 protein ID |  | notes |
| --- | --- | --- | --- | --- | --- | --- |
| *cluster* | *Annotation* | *this study* | *GenBank* | *this study* | *GenBank* |  |
| MCP | Major Capsid Protein | Yara@MT293574_40 | YP_010800661.1 | Ple2@KY346835_31 | AUD57312.1 | core *Nucleocytoviricota* (NCVOG0022) |
| ATPase | DNA packaging ATPase | Yara@MT293574_39 | YP_010800660.1 | Ple2@KY346835_25 | AUD57306.1 | core *Nucleocytoviricota* (NCVOG0249) |
| VLTF3 | virus late transcription factor 3 | Yara@MT293574_51 | YP_010800673.1 | Ple2@KY346835_24 | AUD57305.1 | core *Nucleocytoviricota* (NCVOG0262) |
| RuvC | RuvC-like Holliday junction resolvase | Yara@MT293574_67 | YP_010800690.1 | Ple2@KY346835_33 | AUD57314.1 | core *Nucleocytoviricota* (NCVOG0278) |
| PDDEXK | PDDEXK superfamily endonuclease | Yara@MT293574_2 | YP_010800622.1 | Ple2@KY346835_28 | AUD57309.1 | probable bacterial/phage origin; present in *Nucleocytoviricota* |
| SF3_hel1 | SF3 family helicase | Yara@MT293574_66 | YP_010800689.1 |  |  |  |
| SF3_hel2 | SF3 family helicase |  |  | Ple2@KY346835_2045..6606 | AUD57284.1 |  |
| VLTF2 | virus late transcription factor 2 |  |  | Ple2@KY346835_13 | AUD57294.1 | core *Nucleocytoviricota* (NCVOG1164) |
| Mriya_1 |  | Yara@MT293574_1 | YP_010800621.1 | Ple2@KY346835_29 | AUD57310.1 | Proteins of variable length; only one domain of about 100 amino acids conserved. |
| Mriya_51 |  | Yara@MT293574_50 | YP_010800671.1 | Ple2@KY346835_22 | AUD57304.1 | High count in Yaravirus proteomics |
| Mriya_50 |  | Yara@MT293574_49 | YP_010800670.1 | Ple2@KY346835_21 | AUD57302.1 | Two TM helices; uncharacterized homologs in mimiviruses and phycodnaviruses. |
| Mriya_48 | envelope protein | Yara@MT293574_47 | YP_010800668.1 | Ple2@KY346835_19 | AUD57300.1 | Homologs in some members of *Nucleocytoviricota* (NCVOG1423) |
| HUH | HUH endonuclease | Yara@MT293574_27 | YP_010800648.1 | Ple2@KY346835_11 | AUD57292.1 |  |
| HUH_long | HUH endonuclease domain protein |  |  | Ple2@KY346835_10 | AUD57291.1 |  |
| PEV_8 |  |  |  | Ple2@KY346835_8 | AUD57289.1 |  |
| PEV_12 |  |  |  | Ple2@KY346835_12 | AUD57293.1 |  |
| PEV_14 |  |  |  | Ple2@KY346835_14 | AUD57295.1 |  |
| PEV_15 |  |  |  | Ple2@KY346835_15 | AUD57296.1 |  |
| PEV_20 |  |  |  | Ple2@KY346835_20 | AUD57301.1 |  |
| PEV_22 | minor capsid protein (mCP) | Yara@MT293574_45 | YP_010800666.1 | Ple2@KY346835_23 | AUD57303.1 | high count in Yaravirus proteomics |
| PEV_26 | ssDNA binding protein (ssb) |  |  | Ple2@KY346835_26 | AUD57307.1 |  |
| PEV_27 |  |  |  | Ple2@KY346835_27 | AUD57308.1 |  |
| PEV_34 |  |  |  | Ple2@KY346835_34 | AUD57316.1 |  |
| PEV_35 |  |  |  | Ple2@KY346835_35 | AUD57315.1 |  |

Given the relatively low sequence conservation among the MCPs, we made separate alignments for *Yaraviridae* (Figure S1) and “*Gamadviridae*” (Figure S2) and used each of these MCP alignments as queries to search the PDB, Pfam_A, UniProt-SwissProt-viral, and NCBI_Conserved_Domains (CD) databases using HHPred. This search retrieved, with highly significant scores, the MCP sequences from several major groups in the phylum *Nucleocytovirivota* including members of the families *Mimiviridae, Iridoviridae, Ascoviridae*, and *Phycodnaviridae*, supporting the affiliation of mriyaviruses with *Nucleocytoviricota* (Figure S3). Furthermore, AF2 modeling of the mriyavirus MCP structure followed by comparison with the available diverse structures of DJR MCPs also demonstrated the greater similarity between mriyaviruses and members of the *Nucleocytoviricota* (Figure 4). In the structure-based comparison, *Yaraviridae* and “*Gamadviridae*” formed two separate clades, with *Yaraviridae* showing closer structural similarity to the MCPs of *Nucleocytoviricota*. The position of mriyaviruses between MCPs of polintons and *Nucleocytoviricota,* that is, at the base of the *Nucleocytoviricota* (Figure 4), suggests that, unlike “*Mininucleoviridae*”, mriyaviruses are not diminutive derivatives of *Nucleocytoviricota*, but could rather represent the lineage that, among the currently known viruses, most closely resembles the ancestors of the *Nucleocytoviricota*. Phylogenetic analysis of the packaging ATPase and VLTF3, which are conserved in all mriyaviruses and nearly all members of the *Nucleocytoviricota* (Figure 3), supported the mriyavirus monophyly and was compatible with the basal position of mriyaviruses with respect to *Nucleocytoviricota* (Figure 1b,c).

**Figure 4.**
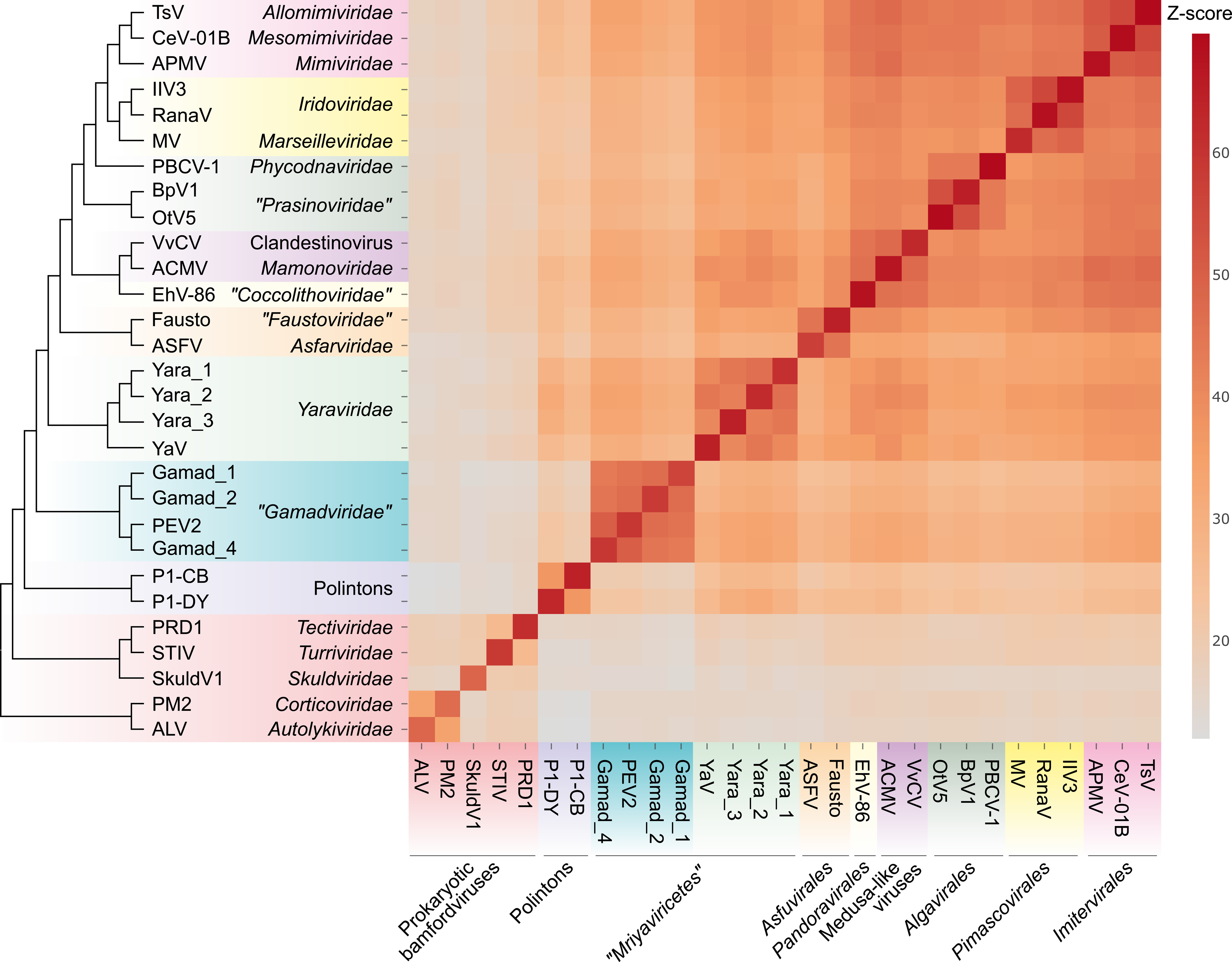
Comparison of the predicted structures of mriyavirus major capsid proteins with structures of major capsid proteins of other members of the kingdom *Bamfordvirae*. The heat map reflects the z-scores obtained in structural comparisons of the MCPs using Dali (color gradient shown to the right of the heat map). The dendrogram shows clustering of the MCPs by the z-scores. The abbreviations are as follows: TsV, *Tetraselmis* virus 1 (YP_010783039); CeV-01B, *Chrysochromulina ericina* virus 01B (YP_009173446); APMV, *Acanthamoeba polyphaga* mimivirus (ADO18196.2); IIV3, Invertebrate iridescent virus 3 (YP_654586); RanaV, Ranavirus maximus (YP_009272725); MV, Marseillevirus marseillevirus (YP_003407071); PBCV-1, *Paramecium bursaria chlorella* virus 1 (PDB id: 5tip); BpV1, Bathycoccus sp. RCC1105 virus BpV1 (YP_004061587); OtV5, *Ostreococcus tauri* virus 5 (YP_001648266); VvCV, V*ermamoeba vermiformis* clandestinovirus (QYA18424); ACMV, *Acanthamoeba castellanii* medusavirus (BBI30317); EhV-86, *Emiliania huxleyi* virus 86 (YP_293839); Fausto, Faustovirus (PDB id: 5j7o); ASFV, African swine fever virus (PDB id: 6ku9); Yara_1, Ga0364485_12008_8; Yara_2, Ga0466970_0005716_5; Yara_3, Yara_group_Contig_26_5; YaV, *Yaravirus brasiliensis* (QKE44414); Gamad_1, Ga0181388_1000587_17; Gamad_2, Ga0314846_0002864_7; PEV2, *Pleurochrysis* sp. endemic virus 2 (AUD57312); Gamad_4, pleuro_group_Assembly_Contig_24_24; P1-CB, Polinton 1 of *Caenorhabditis briggsae*; P1-DY, Polinton 1 of *Drosophila yakuba*; PRD1, *Enterobacteria* phage PRD1 (PDB id:1hx6); STIV, *Sulfolobus* turreted icosahedral virus 1 (PDB id: 3j31); SkuldV1, *Lokiarchaea* virus SkuldV1 (UPO70972); PM2, *Pseudoalteromonas* phage PM2 (PDB id: 2vvf); ALV, *Vibrio* phage 1.020.O._10N.222.48.A2 (AUR82054).

VLTF2 is conserved in all “*Gamadviridae*” and some of the *Yaraviridae*, suggesting that the common ancestor of mriyaviruses encoded this protein. The alignment of VLTF2 protein sequences (Figure S4) contained too few conserved positions to allow reliable tree construction. Nevertheless, HHpred search initiated with the mriyavirus VLTF2 alignment retrieved VLTF2 proteins of different virus families within *Nucleocytoviricota*, in particular, poxviruses (Figure S5), further supporting the link between mriyaviruses and *Nucleocytoviri*cota.

The RuvC-like protein, a homolog of the Holliday junction resolvase encoded by most members of the *Nucleocytoviricota*, is conserved in nearly all mriyaviruses (Figure 3). However, the (predicted) resolvases of mriyaviruses and those of the members of *Nucleocytoviricota*, and even the RuvC-like proteins of different groups of mriyaviruses themselves might be polyphyletic. Indeed, HHpred search initiated from gamadvirus RuvC-like protein sequence alignment retrieved poxvirus RuvC as the top hits (Figure S6) whereas the search initiated with yaravirus RuvC alignment retrieved bacterial and phage homologs first (Figure S6). The alignment of mriyavirus RuvC sequences with their closest homologs included few conserved positions apart from the catalytic motifs, and the phylogenetic tree reconstructed from this alignment was unreliable (Figure S7).

Nearly all (55 out of the 60) representative genomes of mriyaviruses encode helicases of either Superfamily 3 (SF3) or Superfamily 2 (SF2) (Figure 5). The SF3 helicases formed two distinct clusters by sequence similarity: SF3_hel1 represented in most of the members of *Yaraviridae* and Phaglo_G, whereas SF3_hel2 conserved in “*Gamadviridae*”. The sequences of the SF3 helicases were not highly similar to those that are encoded by all members of the *Nucleocytoviricota*, and phylogenetic analysis of the helicases suggested that mriyaviruses have acquired these proteins from bacteriophages or plasmids, independently of the *Nucleocytoviricota* (Figure S9). The SF2 helicases (SF2_hel) were found in a relatively small subset of *Yaraviridae* members (Figure 3) and showed the closest similarity to the mimivirus R8 (AAV50283) and African swine fever virus (ASFV) pF1055L (P0CA09) helicases (15), which are related to the more extensively studied origin-binding protein UL9 conserved in herpesviruses and malacoherpesviruses (16). The helicase domains in all mriyaviruses are the C-terminal regions of larger, apparently multidomain proteins. The N-terminal regions of these proteins are noticeably less conserved than the helicases. This protein architecture resembles one of the universally conserved proteins of the *Nucleocytoviricota* (exemplified by poxvirus D5 protein) that consists of an N-terminal archaeo-eukaryotic primase (AEP) domain and a C-terminal SF3 helicase domain (Figure 5a). However, among the 3 helicase groups, only SF2_hel contained a conserved, intact AEP domain that is also conserved in the homologous proteins of mimiviruses, ASFV and Ostreid herpesvirus 1 (AAS00940; *Malacoherpesviridae*), but not in the UL9-like proteins of mammalian orthoherpesviruses (Fig. 5b,c).

**Figure 5.**
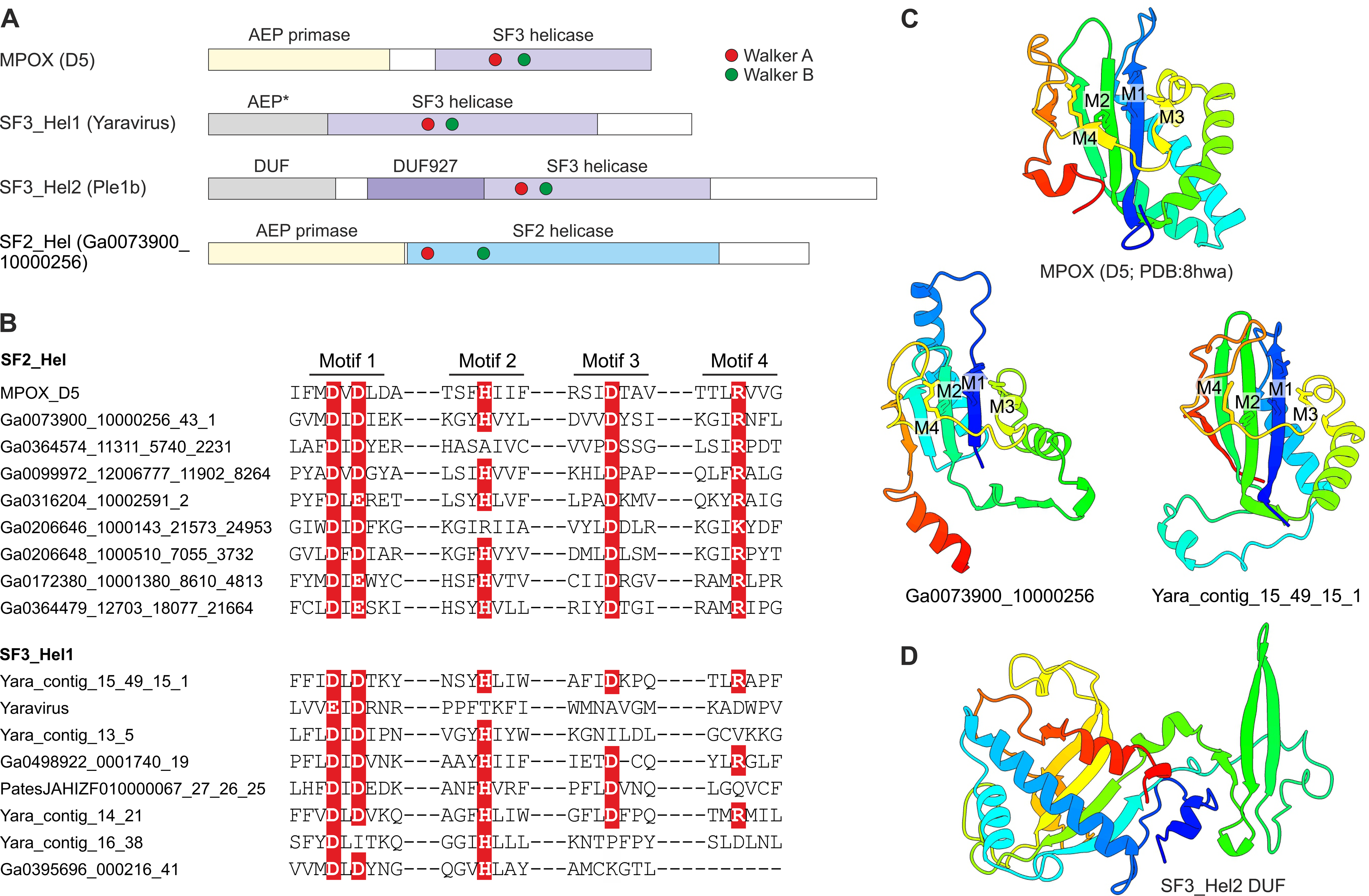
The helicase-containing proteins of mriyaviruses. A, Domain architectures of the helicase-containing proteins of mriyaviruses and the poxvirus primase-helicase (D5) shown for comparison. The asterisk indicates that in the SF3_Hel1 group, most of the AEP homologs contain disrupted catalytic motifs and thus appear to be inactivated. DUF, Domain of Unknown Function; MPOX, Monkeypox virus. B, Sequence segments of AEP catalytic motifs of selected SF2_Hel and SF3_Hel1 proteins. The residues implicated in catalysis are show with white letters on red background. C, Structural models of predicted AEPs of the SF3_Hel1 and SF2_Hel groups of mriyavirus proteins compared to the structure of the AEP domain of MPOX (pdb accession indicated). M1-M4 denote AEP catalytic motifs shown in Figure 5b. D, Structural model of the DUF located at the N-terminus of the SF3_Hel2 proteins.

Despite the considerable divergence within the SF2_hel group, the alignment of these proteins encompassed the four catalytic motifs characteristic of the AEP superfamily primases (17, 18) (Figure 5b), and structure of the AEP domain could be confidently modeled, revealing a characteristic RNA-recognition motif (RRM) (Fig. 5c) (Fig. 5b,c). Notably, the histidine of Motif 2 involved in nucleotide binding is mutated to alanine or arginine in some SF2_Hel proteins (Fig. 5b). However, substitutions within this motif are not uncommon in primases encoded by bacterial and archaeal mobile elements (18), suggesting that the N-terminal domain of mriyavirus SF2_hel is an active primase. In addition, the AEP motifs were detected in the SF3_hel1 proteins (but none of the SF3_hel2) (Figure 5b), and structural modeling supported the similarity to AEP (Figure 5c). However, in most members of SF3_Hel1, some of the catalytic residues of the AEP are replaced (Figure 5b), suggesting that the AEP domain was undergoing degradation during the evolution of the *Yaraviridae*, in most cases, likely losing the primase activity. The N-terminal regions of SF3_hel2 proteins showed no sequence similarity to known domains, and, although a high quality model of this globular domain was obtained using AF2 (Fig. 5d), DALI searches against the PDB database did not reveal any structurally similar domains. Some of the SF3_hel2 genes contain frameshifts in the 5’-terminal region (Figure S10), compatible with the degradation of the N-terminal domain of this protein and suggesting that it is not essential for viral genome replication. Overall, these findings suggest that replication of the mriyavirus genomes requires a DNA helicase; the SF2 and SF3 helicases are mutually exclusive among mriyaviruses, indicating that they are functionally equivalent. By contrast, primase activity is unlikely to be required for mriyavirus replication although the primase domain of the SF2_hel proteins might have an additional function.

Unexpectedly, we found that all mriyaviruses encode a HUH family endonuclease that is involved in the rolling circle replication initiation of the ssDNA viruses of the realm *Monodnaviria* as well as diverse small plasmids and some viruses with dsDNA genomes (19–21). The sequence motifs characteristic of the catalytic site of HUH endonucleases are conserved in all mriyavirus homologs (Figure 6a). The HHpred search initiated with the mriyavirus protein sequences retrieved replication endonucleases of various viruses with highly significant scores (Figure S11). The highest score was obtained with the HUH endonuclease of *Sulofolobus islandicus* rudivirus 1 (SIRV1), a dsDNA virus, and structural analysis yielded a near perfect superposition of the mriyavirus HUH domains with the crystal structure of this protein (Figure 6b), with the predicted catalytic amino acid residues juxtaposed to form the catalytic site (Figure 6c). These findings strongly suggest that all mriyaviruses encode an active rolling circle replication initiation endonuclease. Gamadviruses additionally encode a larger protein conserved within this group that contains a C-terminal HUH endonuclease domain and an uncharacterized N-terminal region (Figure 3). The two HUH domains of gamadviruses are closely similar (Figure 6a,d) suggesting a duplication at the onset of gamadvirus evolution followed by the capture of the additional N-terminal domain. The conservation of the HUH endonuclease and its catalytic motifs in all mriyaviruses strongly suggests that the endonuclease activity of this protein is essential for replication.

**Figure 6.**
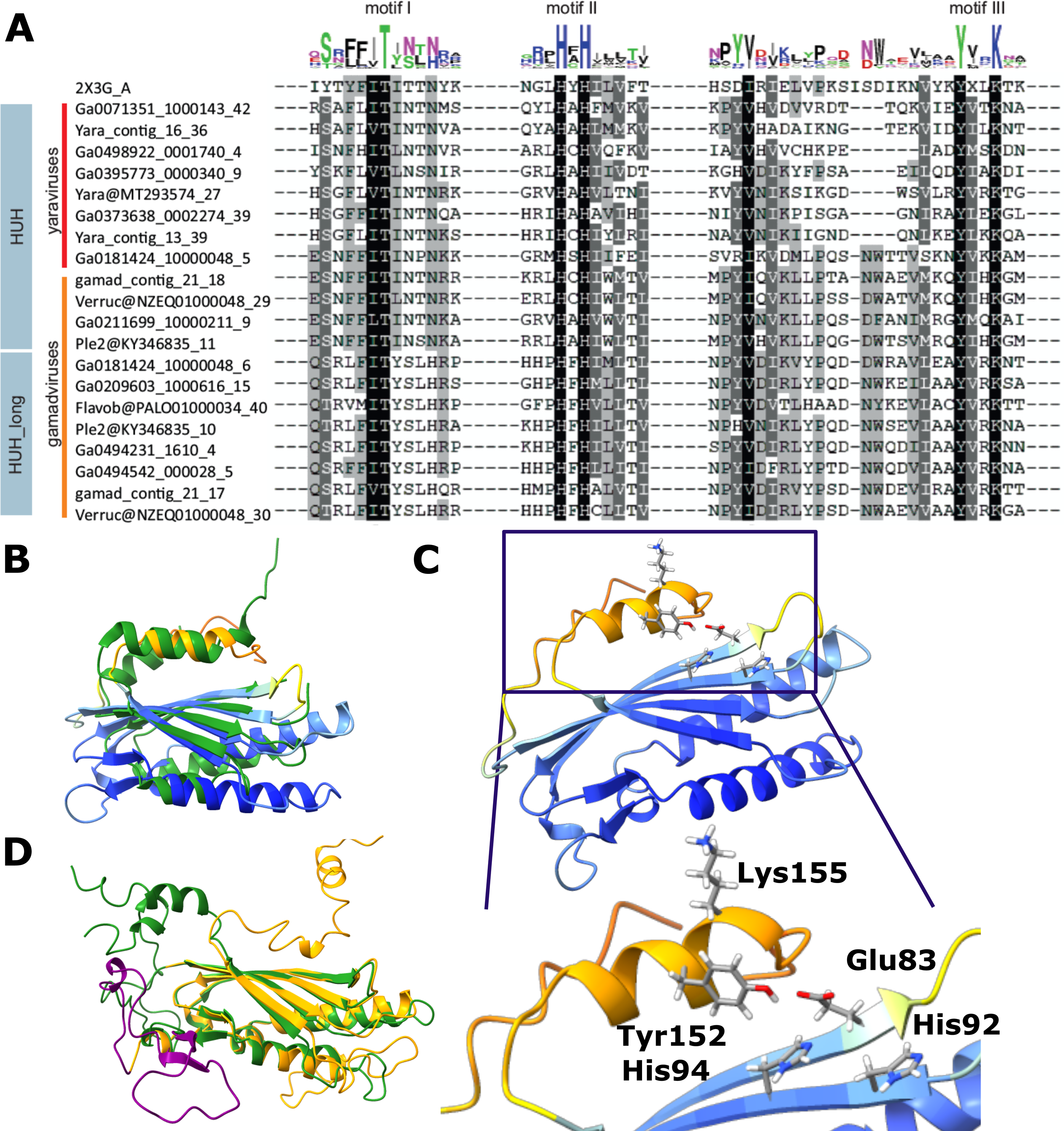
Sequence and structure conservation in the HUH endonucleases of mriyaviruses. A, Alignment of the sequence segments of the HUH superfamily endonucleases containing the characteristic motifs I-III (N-terminal motif I consisting of hydrophobic residues, motif II with (HUH; H: Histidine, U: hydrophobic residue) and C-terminal motif III (Yx2-3K; Y: tyrosine, x: any residue, K: lysine, blue), where only the second tyrosine is present (compared to the full motif 3 YUxxYx2-3K, U: hydrophobic residue), are highlighted. B, A representative predicted structure of a mriyavirus HUH endonuclease superimposed with the crystal structure of protein ORF119 from *Sulfolobus islandicus* rod-shaped virus 1 (green, pdb 2X3G-A, z-score 7.7). Yaravirus HUH endonuclease (MT293574_27) colored by plddt score. C, Configuration of the catalytic amino acid residues of motif II and III in the predicted structure of the mriyavirus HUH endonuclease (Yaravirus MT293574_27, colored by plddt score). D, Superposition of the structural models of the two HUH endonuclease domains of gamadviruses (short, probably active: KY346835_11 (green, aa 31-224, aa1-30 unstructured, clipped off for representation), long: KY346835_10 (orange, aa 1353-1574 with additional inserted loop (purple) aa 104-1450).

Most Mriyaviruses encode a PDDEXK superfamily endonuclease (Figure 3 and Figure S12). This protein is homologous but apparently not orthologous to the viral recombinase YqaJ that is encoded by many members of the *Nucleocytoviricota* (2). Rather, the mriyavirus PDDEXK endonuclease is likely to be of bacterial or phage origin as indicated by the phylogenetic tree topology (Figure S13).

Yaravirus gene 48 (numbered as in Boratto et al., 2020) encodes a protein that is conserved in mriyaviruses and for which homologs with significant sequence similarity were detected in many members of *Nucleocytoviricota* including mimiviruses, phycodnaviruses and iridoviruses as well as other viruses and bacteria (Figure S14). The phylogenetic tree of these proteins (hereafter Mriya_48) is compatible with the monophyly of mriyaviruses but does not imply a direct connection to *Nucleocytoviricota (*Figure S15a). The iridovirus homologs of Mriya_48 are structural proteins located in the virion envelope (22). Structural comparison of Mriya_48 and the iridovirus envelope protein ORF056L (GenBank ID: NP_612278) revealed pronounced structural similarity, suggestive of similar functions (Figure 7). However, the predicted structure of the gene 48 product of Yaravirus itself failed to superimpose with the iridovirus envelope protein due to the apparent different spatial arrangements of the α-helices (Figure S16b). Considering that this protein was not detected in the Yaravirus particle proteome (12), it might have lost its function as an envelope protein in Yaravirus.

**Figure 7.**
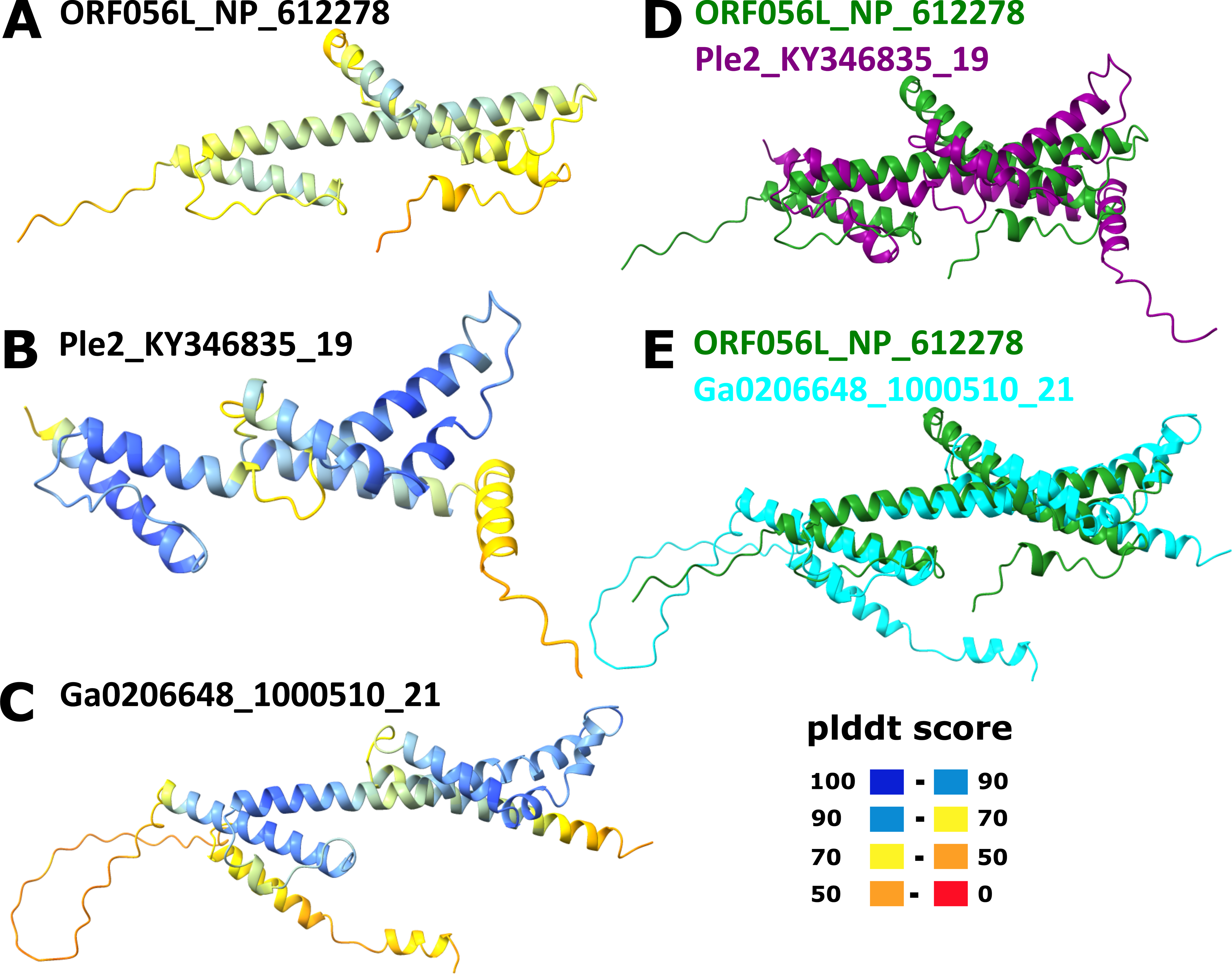
Comparison of the structural models of Mriya_48 protein and iridovirus envelope protein. A, Iridovirus enveloped protein (ORF056L_NP_612278); B, Ple2_KY346835_19; C, Ga0206648_1000510_21; D, Superposition of ORF056L_NP_612278 (green) and Ple2_KY346835_19 (purple); E, Superposition of ORF056L_NP_612278 (green) and Ga0206648_1000510_21 (cyan). In A-C, the structures are colored according to the plddt score.

The rest of the proteins conserved across the mriyaviruses either lacked detectable homologs outside this group of viruses or at least lacked functionally characterized homologs (Table 1 and Figures S16-S20). In addition to the 12 proteins comprising the Mriyavirus core, 10 more proteins were found to be conserved in members of the “*Gamadviridae*” (Figure 3 and Supplementary Table S1). One of these proteins, PEV_22 (numbered as in PEV 2), was identified as the minor capsid protein containing a typical single jelly roll domain and structurally similar to the minor capsid protein of Mavirus virophage (Figure S21). Another protein conserved in gamadviruses is PEV_26, which is predicted to be structurally similar to the OB-fold containing single-stranded DNA-binding (SSB) protein of bacteriophage T7 (PDB structure 1je5; Figure S22). Putative SSB homologous to the T7 SSB have been previously identified in 4 virus families within *Nucleocytoviricota* (*Phycodnaviridae*, *Mimiviridae*, *Iridoviridae* and *Marseilleviridae*) (23). The remaining 8 proteins conserved in “*Gamadviridae*” remain uncharacterized, without detectable homologs.

The identification of a candidate minor capsid protein in gamadviruses prompted us to search for a counterpart in the members of *Yaraviridae*. We found that the product of Yaravirus gene 46 (YP_010800666), the second most abundant protein in the Yaravirus virion proteome, contains a predicted single jelly roll domain at its C-terminus (Figure S23) and thus is a strong candidate for the minor capsid protein. Indeed, a PSI-BLAST search initiated with this protein sequence retrieved uncharacterized proteins of some members of the *Nucleocytoviricota* (marseilleviruses, medusaviruses) as well as Sputnik and Zamilon virophage minor capsid proteins (Figure S24). The uncharacterized homologous proteins in *Nucleocytoviricota* had the same modular architecture as Yaravirus gene 46 consisting of a C-terminal single jelly-roll domain (TNF superfamily) and a variable N-terminal domain that is predicted to adopt either an α-helical or a β-sheet fold; in some of these proteins, the N-terminal domain appears to be disordered or is missing altogether. In contrast, the Sputnik and Zamilon virophage minor capsid proteins consist of two domains, a ‘lower’ single jelly-roll domain and an ‘upper’ β-barrel domain inserted between β-strands D and E of the jelly-roll (24). Modeling and comparing all proteins of 35 representatives of *Yaraviridae* led to the detection of a Sputnik penton-like minor capsid protein encoded in 22 genomes (with a likely duplication in Ga0209319) whereas the single jelly-roll C-terminal domain with the variable N-terminus was less common (7 genomes, with 3 paralogs in Yaravirus MT293574 (genes 11, 12 and 45). Only 2 yaravirus genomes (Ga0172380_10001380 and Ga0182030_10004970) were found to encode both types of putative minor capsid proteins. Thus, in accord with previous observations on *Nucleocytoviricota* (25), the minor capsid proteins of mriyaviruses appear to be highly variable and candidates for this role remain to be identified in some member of “*Mriyaviricetes*”.

## Discussion

In this work, by mining genomic and metagenomic databases, we identified a distinct group of viruses that appear to be related to the members of the phylum *Nucleocytoviricota* but have genomes in the 35-45 kb range, much smaller than the genomes of any previously known members of this phylum. We coined the name Mriyaviruses for this group. Mriyaviruses include the previously identified Yaravirus and PEV as well as about 200 apparently complete or near-complete viral genomes identified in metagenomes. Yaravirus was isolated by cultivation in *Acanthamoeba castellanii* (12) whereas PEV infect a haptophyte host. The related viruses identified in this work come from metagenomes representing a broad diversity of environments suggesting that mriyaviruses infect diverse unicellular eukaryotes.

The majority of the proteins encoded by mriyaviruses (>90% as reported in the original analysis of the Yaravirus genome (12)) showed no readily detectable sequence similarity to any known proteins. Nevertheless, through a combination of sensitive sequence searches with protein structure modeling followed by search of structural databases for potential homologs, we established the identity of many mriyavirus gene products. Five of these proteins are also conserved among most members of the phylum *Nucleocytoviricota* and two more had homologs within more limited subsets of the phylum members (Table 1) enabling phylogenetic analysis and evolutionary inferences. The evolutionary provenance of mriyaviruses did not appear to be immediately obvious given that, in terms of the genome size, they are closer to the viruses of the phylum *Preplasmiviricota* (such as polintons, adenoviruses or virophages) that, together with the phylum *Nucleocytoviricota*, belongs to the kingdom *Bamfordvirae* within the realm *Varidnaviria* and shares with the latter the homologous MCP, minor capsid protein and packaging ATPase (26). Nevertheless, the presence of two signature genes of *Nucleocytoviricota*, VLTF2 and VLTF3, along with the results of structural comparisons of the MCPs, strongly suggests that Mriyaviruses are a distinct branch of *Nucleocytoviricota*. Phylogenetic analysis and structural comparison of the conserved proteins does not point to an affinity between mriyaviruses and any particular clade of *Nucleocytoviricota*, suggesting that these viruses should be assigned the rank of class, *“Mriyaviricetes”*. In the phylogenies, “*Mriyaviricetes*” split into two distinct clades, one of which is a compact group including viruses related to PEV, for which we propose the name “*Gamadviridae*” (possibly, to be elevated to the order rank) and the other one is a looser group corresponding to the family *Yaraviridae* (possibly, another order in the future).

Mriyaviruses encode no RNA polymerase subunits suggesting that, similarly to “*Mininucleoviridae*” (11), they reproduce in the nuclei of the host cells. Mriyaviruses encode a small but unusual set of proteins implicated in viral genome replication. As noticed also for other large dsDNA viruses (27, 28), the replication machinery components are not strongly conserved among the members of “*Mriyaviricetes*”, with several ancestral genes apparently replaced with genes of different origins encoding proteins with the same functions. Mriyaviruses lack the DNA-dependent DNA polymerase that is encoded by all other members of the *Nucleocytoviricota*, with the obvious implication that the replication of mriyavirus genomes relies on a host DNA polymerase. Almost all mriyaviruses encode helicases (either SF3 or SF2) that in some cases are fused to primase (AEP) domains, which is another signature of the *Nucleocytoviricota* (2). However, the AEP is predicted to be active only in a minority of the mriyaviruses, whereas the majority contain either an AEP that appears to be inactivated or an uncharacterized N-terminal domain. Unexpectedly, all mriyaviruses were found to encode an HUH superfamily endonuclease (duplicated in “*Gamadviridae*”), the enzyme that is involved in the initiation of rolling circle replication of the ssDNA viruses of the realm *Monodnaviria*, diverse small plasmids and some dsDNA viruses (20). In particular, HUH endonucleases are also encoded by varidnaviruses of at least two families, *Corticoviridae* (29, 30) and *Simuloviridae* (31, 32), in which they apparently were acquired independently (21). The combination of primase and HUH endonuclease, proteins associated with different modes of genome replication, to our knowledge, so far has not been observed in any viruses or plasmids (27). By contrast, eukaryotic monodnaviruses typically encode both an HUH endonuclease and a SF3 helicase as a fusion protein (21).

Analysis of the proteins implicated in mriyavirus genome replication allows us to propose a plausible evolutionary scenario. Given that AEP is conserved and appears to be essential for genome replication in all members of the *Nucleocytoviricota* (2), it seems likely that the ancestral mriyavirus replicated via the same, RNA-primed mechanism. However, subsequent acquisition of the HUH endonuclease, which is conserved and predicted to be active in all mriyaviruses, suggests that this protein is essential for replication and is likely to initiate replication via a rolling circle mechanism as demonstrated for P2-like bacteriophages that have similar-sized, 33 kb genomes (33–35). The switch of the replication mode in mriyaviruses apparently was accompanied by the loss of the primase activity or apparent replacement of the AEP domain with an unrelated domain in different lineages of mriyaviruses. The helicase, in contrast, was retained or replaced by a distinct one, at least, in most mriyaviruses, and likely interacts with the HUH endonuclease during replication. Indeed, whereas eukaryotic HUH endonucleases function with the cognate SF3 helicases, bacteriophages that replicate by rolling circle mechanism hijack host SF1 helicases (36, 37), further suggesting that DNA unwinding during the rolling circle replication can be carried out by a broad variety of helicases.

Perhaps, the most intriguing feature of mriyaviruses is their putative ancestral status with respect to the rest of the member of the *Nucleocytoviricota* as indicated by their deep placement in phylogenetic trees of the conserved proteins and by comparison of the MCP structures. Further expansion of the “*Mriyaviricetes*” through extended metagenome mining and/or discovery of additional groups of viruses with small genomes related to the *Nucleocytoviricota* can be expected to further clarify and solidify the scenario for the origin and evolution of this expansive phylum of bamfordviruses.

## Conclusions

In this work, we describe a distinct group of dsDNA viruses, mriyaviruses, that share 5 conserved genes with large and giant viruses of the phylum *Nucleocytoviricota* and, based on this commonality and structural comparisons of the MCPs, appear to belong to this phylum although they have comparatively small genomes of only 35-45 kb. The previously characterized mriyaviruses, Yaravirus and PEV, infect amoeba and haptophytes, respectively, and the genomes of other mriyaviruses were assembled from metagenomes originating from a variety of environments, suggesting that mriyaviruses infect diverse unicellular eukaryotes. Phylogenetic analysis does not reveal specific affinity between mriyaviruses and any other branch of the *Nucleocytoviricota*, suggesting that these viruses comprise a separate class, “*Mriyaviricetes*”. Structural comparisons of the MCPs suggest that mriyaviruses could be the lineage that, among the known groups of viruses, is most closely related to the ancestors of the *Nucleocytoviricota*. In phylogenetic trees, mriyaviruses split into two well-separated branches, the family *Yaraviridae* and proposed family “*Gamadviridae*”. Mriyaviruses lack DNA polymerase which is encoded by all other members of the *Nucleocytoviricota* and RNA polymerase subunits encoded by all members of the *Nucleocytoviricota* that reproduce in the host cell cytoplasm. Thus, mriyaviruses probably replicate in the host cell nuclei. Mriyaviruses encode both a helicase-primase, which is an essential component of the DNA replication apparatus of the *Nucleocytoviricota,* and a HUH endonuclease, a combination so far not found in any viruses. The primase domain is inactivated or replaced in most mriyaviruses whereas the HUH endonuclease is conserved and predicted to be active in all members of the “*Mriyaviricetes*”, suggesting that its activity is essential for the initiation of mriyavirus genome replication via the rolling circle mechanism.

## Materials and Methods

### Collecting mriyavirus MCP-encoding contigs

Publicly available genomic (NCBI GenBank; https://www.ncbi.nlm.nih.gov/genbank) and metagenomic (IMG/VR; https://img.jgi.doe.gov/vr) sequence databases were searched using BLASTP (38) for proteins with significant similarity to the MCPs of “*Mininucleoviridae*” (*Panulirus argus* virus 1, GenBank ID QIQ08629.1; *Carcinus maenas* virus 1, QIQ08561.1; *Dikerogammarus haemobaphes* virus 1, QIQ08620.1), *Yaravirus brasiliensis* (YP_010800661.1), and NDDV (Pleurochrysis sp. endemic virus 1a, AUD57260.1; Pleurochrysis sp. endemic virus 1b; AUL80795.1; Pleurochrysis sp. endemic virus 2, AUD57312.1; Pharex and Phaglo_G (Roitman et al., 2023). Genomic sequences encoding proteins with significant similarity to the MCP queries were downloaded and translated using Prodigal in the metagenome mode (39). The predicted proteins were used as queries for a new round of BLASTP search. The retrieved protein sequences were clustered using MMSEQS2 (40), and cluster representatives were aligned with MCPs of representatives of the major groups of *Nucleocytoviricota* (41) using MUSCLE 5 (42). The resulting multiple alignment was used to construct a phylogenetic tree using Fasttree with WAG evolutionary model and Gamma-distributed site rates (43). Based on the MCP tree, mriyavirus MCP-containing contigs were retrieved; several contigs were extended with Geneious Prime® 2022.1.1 (www.geneious.com), to obtain more complete genome sequences (Supplementary Table S1).

### Gene composition and protein function prediction for selected members of “Mriyaviricetes”

A set of 60 genome sequences was selected to represent the mriyavirus sequence diversity (Supplementary Table S1). ORFs were predicted in contigs using Prodigal in the metagenomic mode. Amino acid sequences were initially clustered using MMSEQS2 with the similarity threshold 0.5; the resulting protein clusters were aligned using MUSCLE 5 and iteratively compared to each other using HHSEARCH (44). Clusters of similar sequences (alignment footprint coverage threshold 0.5; relative sequence similarity threshold 0.05) were progressively aligned to each other using HHALIGN (44). The cluster alignments were compared to publicly available profile databases (PDB_mmCIF70, Pfam-A_v36, Uniprot-SwissProt-viral70_3, and NCBI_Conserved_Domains (CD)_v3.19) using HHPRED (for protein annotations, see Supplementary Table S1). Alignment of conserved proteins are available at https://ftp.ncbi.nlm.nih.gov/pub/yutinn/mriya_2024.

### Phylogenetic analysis of conserved proteins of mriyaviruses

A consensus sequence generated from each mriyavirus conserved protein cluster was used as a query to search GenBank (clustered_nr database (45)) for homologous proteins, which were then aligned with mriyavirus proteins using MUSCLE 5. A phylogenetic tree was constructed from this alignment using Fasttree with a WAG evolutionary model and Gamma-distributed site rates. Phylogenetic trees of MCP, packaging ATPase (ATPase), and viral late transcription factor 3 (VLTF3) were built using IQ-TREE (46), with the following models chosen according to BIC by the built-in model finder: Q.pfam+F+R4 for MCP, Q.pfam+F+R6 for ATPase, and VT+F+R5 for VLTF3.

### Protein structure prediction and analysis

Protein structures were modeled using a singularity version of AlphaFold2 version 2.3.2 (47), with the following parameters: “--db_preset=full_dbs –model_preset=monomer_ptm – max_template_date=2023-09-01”) on the high-performance cluster BIOWULF at the NIH. In addition, selected mriyavirus proteins were added to the default uniref90.fasta protein selection of AlphaFold2 (https://ftp.ncbi.nih.gov/pub/yutinn/mriya_2024/mriyavirus_proteins_uniref90.fasta) to improve the quality of alignments generated by AlphaFold2 during its hhsearch run against uniref90. Selected major capsid proteins outside “*Mriyaviricetes*” used for the analysis presented in Figure 4 and not available at pdb were modeled with Colabfold using Alphafold2 multimer v3 (48). Structures were searched against a local version of pdb70 structure database (created 10th of December 2021) using Dali version 5.1 (49) In addition, Foldseek (50) was used to search predicted structures against the Foldseek databases ‘AlphaFold proteome’, ‘AlphaFold swissprot’ (both version 2) and ‘pdb’ (version from 2023-08-20). Comparison of predicted and experimentally resolved structures from pdb for selected mriyavirus and *Nucleocytoviricota* major capsid protein homologs was performed by running Dali all-vs-all. Protein structures and structural models were visualized using Chimera X (51).

#### Data availability

This paper is based entirely on the analysis of existing, publicly available data. Data generated during downstream analysis are available in the Supplementary Material or via ftp at https://ftp.ncbi.nih.gov/pub/yutinn/mriya_2024. Any additional information required to reanalyze the data reported in this paper is available from the authors.

## Author contributions

N.Y. and E.V.K. initiated the study; N.Y. collected the data; N.Y., P.M., M.K. and E.V.K. analyzed the data; N.Y. and E.V.K. wrote the manuscript that was edited and approved by all authors.

## Acknowledgements

The authors thank Darius Kazlauskas and the Koonin group members for helpful discussions. N.Y., P.M. and E.V.K. are supported through the Intramural Research Program of the National Institutes of Health (National Library of Medicine). This work utilized the computational resources of the NIH HPC Biowulf cluster (http://hpc.nih.gov).

## Notes

### Competing Interest Statement

The authors have declared no competing interest.

## References

1. Abergel C, Legendre M, Claverie JM. 2015. The rapidly expanding universe of giant viruses: Mimivirus, Pandoravirus, Pithovirus and Mollivirus. FEMS Microbiol Rev. 39(6):779–796.

2. Koonin EV, Yutin N. 2019. Evolution of the Large Nucleocytoplasmic DNA Viruses of Eukaryotes and Convergent Origins of Viral Gigantism. Adv Virus Res. 103:167–202.

3. Raoult D, et al. 2004. The 1.2-megabase genome sequence of Mimivirus. Science. 306(5700):1344–1350.

4. Claverie JM, et al. 2009. Mimivirus and Mimiviridae: giant viruses with an increasing number of potential hosts, including corals and sponges. J Invertebr Pathol. 101(3):172–180.

5. Nasir A, Kim KM, Caetano-Anolles G. 2012. Giant viruses coexisted with the cellular ancestors and represent a distinct supergroup along with superkingdoms Archaea, Bacteria and Eukarya. BMC Evol Biol. 12:156.

6. Nasir A, Kim KM, Caetano-Anolles G. 2017. Phylogenetic Tracings of Proteome Size Support the Gradual Accretion of Protein Structural Domains and the Early Origin of Viruses from Primordial Cells. Front Microbiol. 8:1178.

7. Moreira D, Lopez-Garcia P. 2005. Comment on “The 1.2-megabase genome sequence of Mimivirus”. Science. 308(5725):1114; author reply 1114.

8. Williams TA, Embley TM, Heinz E. 2011. Informational gene phylogenies do not support a fourth domain of life for nucleocytoplasmic large DNA viruses. PLoS One. 6(6):e21080.

9. Yutin N, Wolf YI, Koonin EV. 2014. Origin of giant viruses from smaller DNA viruses not from a fourth domain of cellular life. Virology.

10. Koonin EV, Yutin N. 2018. Multiple evolutionary origins of giant viruses. F1000Res. 7.

11. Subramaniam K, et al. 2020. A New Family of DNA Viruses Causing Disease in Crustaceans from Diverse Aquatic Biomes. mBio. 11(1).

12. Boratto PVM, et al. 2020. Yaravirus: A novel 80-nm virus infecting Acanthamoeba castellanii. Proc Natl Acad Sci U S A. 117(28):16579–16586.

13. de Miranda Boratto PV, Oliveira GP, Abrahao JS. 2022. “Yaraviridae”: a proposed new family of viruses infecting Acanthamoeba castellanii. Arch Virol. 167(2):711–715.

14. Roitman S, et al. 2023. Isolation and infection cycle of a polinton-like virus virophage in an abundant marine alga. Nat Microbiol. 8(2):332–346.

15. Gupta A, Patil S, Vijayakumar R, Kondabagil K. 2017. The Polyphyletic Origins of Primase-Helicase Bifunctional Proteins. J Mol Evol. 85(5-6):188–204.

16. Weller SK, Coen DM. 2012. Herpes simplex viruses: mechanisms of DNA replication. Cold Spring Harb Perspect Biol. 4(9):a013011.

17. Iyer LM, Koonin EV, Leipe DD, Aravind L. 2005. Origin and evolution of the archaeo-eukaryotic primase superfamily and related palm-domain proteins: structural insights and new members. Nucleic Acids Res. 33(12):3875–3896.

18. Kazlauskas D, et al. 2018. Novel Families of Archaeo-Eukaryotic Primases Associated with Mobile Genetic Elements of Bacteria and Archaea. J Mol Biol. 430(5):737–750.

19. Ilyina TV, Koonin EV. 1992. Conserved sequence motifs in the initiator proteins for rolling circle DNA replication encoded by diverse replicons from eubacteria, eucaryotes and archaebacteria. Nucleic Acids Res. 20(13):3279–3285.

20. Chandler M, et al. 2013. Breaking and joining single-stranded DNA: the HUH endonuclease superfamily. Nat Rev Microbiol. 11(8):525–538.

21. Kazlauskas D, Varsani A, Koonin EV, Krupovic M. 2019. Natural history of the rolling-circle replicons: Multiple origins of prokaryotic and eukaryotic ssDNA viruses from bacterial plasmids. Nature Communications. 10(1):3425.

22. Dong CF, et al. 2011. Global landscape of structural proteins of infectious spleen and kidney necrosis virus. J Virol. 85(6):2869–2877.

23. Kazlauskas D, Venclovas C. 2012. Two distinct SSB protein families in nucleo-cytoplasmic large DNA viruses. Bioinformatics. 28(24):3186–3190.

24. Zhang X, et al. 2012. Structure of Sputnik, a virophage, at 3.5-A resolution. Proc Natl Acad Sci U S A. 109(45):18431–18436.

25. Krupovic M, Koonin EV. 2015. Polintons: a hotbed of eukaryotic virus, transposon and plasmid evolution. Nat Rev Microbiol. 13(2):105–115.

26. Koonin EV, et al. 2020. Global Organization and Proposed Megataxonomy of the Virus World. Microbiol Mol Biol Rev. 84(2).

27. Kazlauskas D, Krupovic M, Venclovas C. 2016. The logic of DNA replication in double-stranded DNA viruses: insights from global analysis of viral genomes. Nucleic Acids Res. 44(10):4551–4564.

28. Yutin N, et al. 2021. Analysis of metagenome-assembled viral genomes from the human gut reveals diverse putative CrAss-like phages with unique genomic features Nature Communications. in press.

29. Mannisto RH, Kivela HM, Paulin L, Bamford DH, Bamford JK. 1999. The complete genome sequence of PM2, the first lipid-containing bacterial virus To Be isolated. Virology. 262(2):355–363.

30. Espejo RT, Canelo ES, Sinsheimer RL. 1971. Replication of bacteriophage PM2 deoxyribonucleic acid: a closed circular double-stranded molecule. J Mol Biol. 56(3):597–621.

31. Wang Y, et al. 2018. Rolling-circle replication initiation protein of haloarchaeal sphaerolipovirus SNJ1 is homologous to bacterial transposases of the IS91 family insertion sequences. J Gen Virol. 99(3):416–421.

32. Wang Y, et al. 2016. Identification, Characterization, and Application of the Replicon Region of the Halophilic Temperate Sphaerolipovirus SNJ1. J Bacteriol. 198(14):1952-1964.

33. Odegrip R, Haggard-Ljungquist E. 2001. The two active-site tyrosine residues of the a protein play non-equivalent roles during initiation of rolling circle replication of bacteriophage p2. J Mol Biol. 308(2):147–163.

34. Esposito D, et al. 1996. The complete nucleotide sequence of bacteriophage HP1 DNA. Nucleic Acids Res. 24(12):2360–2368.

35. Sivaprasad AV, Jarvinen R, Puspurs A, Egan JB. 1990. DNA replication studies with coliphage 186. III. A single phage gene is required for phage 186 replication. J Mol Biol. 213(3):449-463.

36. Scott JF, Eisenberg S, Bertsch LL, Kornberg A. 1977. A mechanism of duplex DNA replication revealed by enzymatic studies of phage phi X174: catalytic strand separation in advance of replication. Proc Natl Acad Sci U S A. 74(1):193–197.

37. Arai N, Kornberg A. 1981. Rep protein as a helicase in an active, isolatable replication fork of duplex phi X174 DNA. J Biol Chem. 256(10):5294–5298.

38. Altschul SF, et al. 1997. Gapped BLAST and PSI-BLAST: a new generation of protein database search programs. Nucleic Acids Res. 25(17):3389–3402.

39. Hyatt D, et al. 2010. Prodigal: prokaryotic gene recognition and translation initiation site identification. BMC Bioinformatics. 11:119.

40. Steinegger M, Soding J. 2017. MMseqs2 enables sensitive protein sequence searching for the analysis of massive data sets. Nat Biotechnol. 35(11):1026–1028.

41. Aylward FO, Moniruzzaman M, Ha AD, Koonin EV. 2021. A phylogenomic framework for charting the diversity and evolution of giant viruses. PLoS Biol. 19(10):e3001430.

42. Edgar RC. 2022. Muscle5: High-accuracy alignment ensembles enable unbiased assessments of sequence homology and phylogeny. Nat Commun. 13(1):6968.

43. Price MN, Dehal PS, Arkin AP. 2010. FastTree 2--approximately maximum-likelihood trees for large alignments. PLoS ONE. 5(3):e9490.

44. Soding J. 2005. Protein homology detection by HMM-HMM comparison. Bioinformatics. 21(7):951–960.

45. Sayers EW, et al. 2023. Database resources of the National Center for Biotechnology Information in 2023. Nucleic Acids Res. 51(D1):D29–D38.

46. Minh BQ, et al. 2020. IQ-TREE 2: New Models and Efficient Methods for Phylogenetic Inference in the Genomic Era. Mol Biol Evol. 37(5):1530–1534.

47. Jumper J, et al. 2021. Highly accurate protein structure prediction with AlphaFold. Nature. 596(7873):583–589.

48. Mirdita M, et al. 2022. ColabFold: making protein folding accessible to all. Nat Methods. 19(6):679–682.

49. Holm L. 2020. DALI and the persistence of protein shape. Protein Sci. 29(1):128–140.

50. van Kempen M, et al. 2023. Fast and accurate protein structure search with Foldseek. Nat Biotechnol.

51. Pettersen EF, et al. 2021. UCSF ChimeraX: Structure visualization for researchers, educators, and developers. Protein Sci. 30(1):70–82.

