## Supplementary figures for "Mriyaviruses: Small Relatives of Giant Viruses"

S1 Multiple sequence alignment (MSA) of *Yaraviridae* MCP proteins.

S2 MSA of "*Gamadviridae*" MCP proteins.

S3 Top hits of HHpred searches initiated with *Yaraviridae* and "*Gamadviridae*" MCP protein MSAs.

S4 VLTF2 protein MSA.

S5. Top hits of HHpred searches initiated with Mriyavirus VLTF2 protein MSA.

S6. Top HHpred hits for "*Gamadviridae*" (right) and *Yaraviridae* (left) RuvC MSAs.

S7. Phylogenetic tree of RuvC proteins.

S8. MSA of selected SF3_hel1 proteins.

S9. Phylogenetic tree of SF3_hel1 proteins.

S10. SF3_hel2 protein MSA.

S11. Top hits of HHpred search initiated with Mriyavirus HUH endonuclease MSA.

S12. MSA of HUH and C-terminal domain of HUH_long proteins.

S12. MSA of mriyavirus PDDEXK nuclease.

S13. Phylogenetic tree of PDDEXK nucleases.

S14. MSA of Yaravirus gene 48 and its homologs in mriyaviruses (Mriya_48) and in NCBI Genbank (nr) database.

S15. A, Phylogenetic tree of Mriya_48; B, predicted structures of two Mriya_48 proteins.

S16. MSA of Yaravirus gene 50 and its homologs in mriyaviruses (Mriya_50) and in NCBI Genbank (nr) database.

S17. Predicted structure of Mriya_50 (exemplified with Ple2@KY346835_21). Leucine site chains highlighted in magenta.

S18. MSA of conserved domain of Yaravirus gene 1 and its homologs in mriyaviruses (Mriya_1).

S19. Predicted structure of Mriya_1, exemplified with Ple2@KY346835_29.

S20. A, MSA of Yaravirus gene 51 with mriyavirus and nr homologs. B, Predicted structure of Yaravirus gene 51.

S21. A, Predicted structure of PEV_22; B, same structure overlayed with 6g41_A – the structure of minor capsid protein of Cafeteria mavirus.

S22. Predicted structure of PEV_26 superimposed with gp2.5, a single strand DNA binding protein from bacteriophage T7 (pdb 1je5, green, z-score 13.3).

S23. Predicted structure of Yaravirus minor capsid protein

S24. MSA of Yaravirus minor capsid protein and related sequences.

**S1**

#### *Yaraviridae* MCP

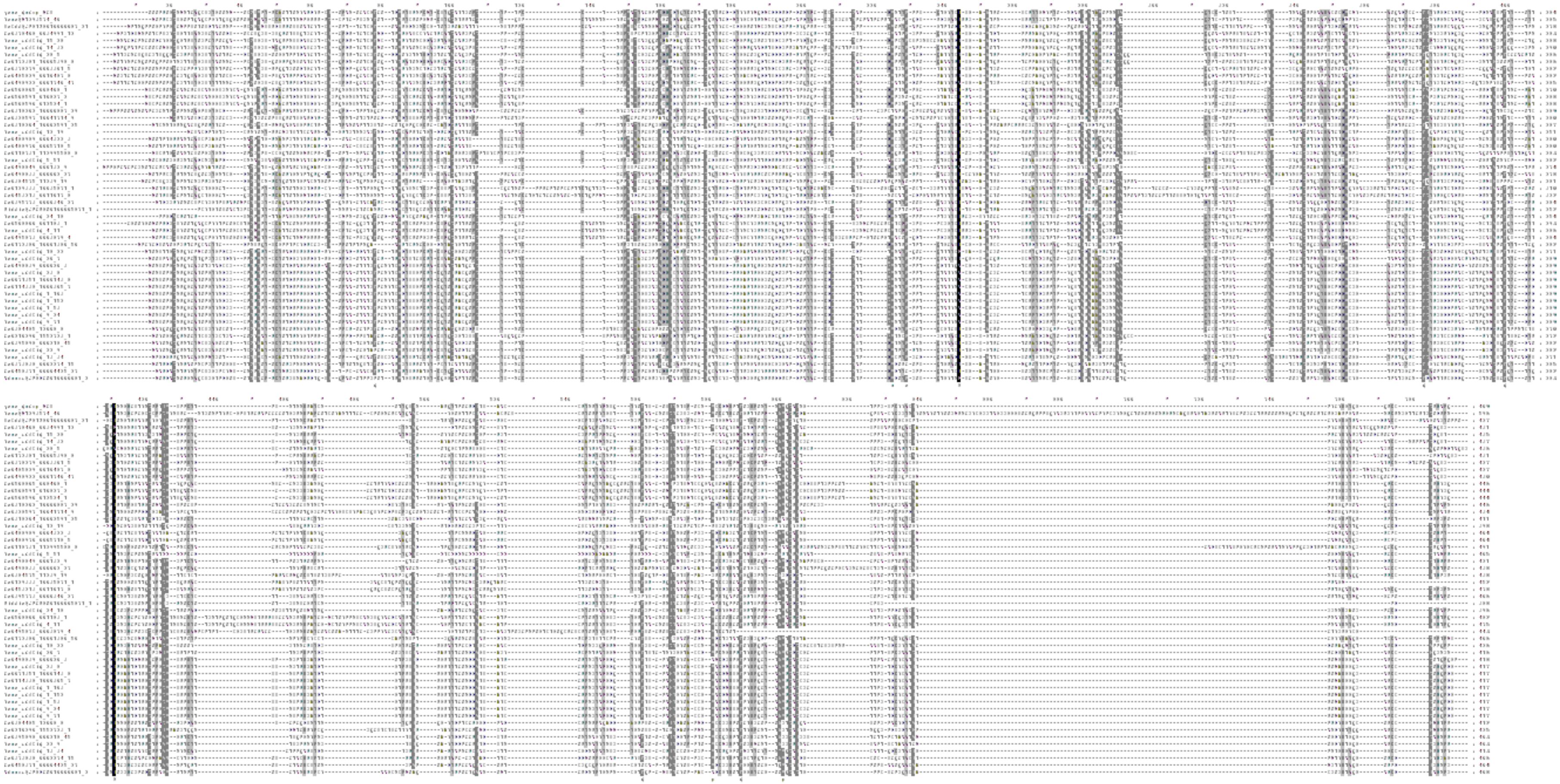

# S2

#### *“Gamadviridae”* MCP

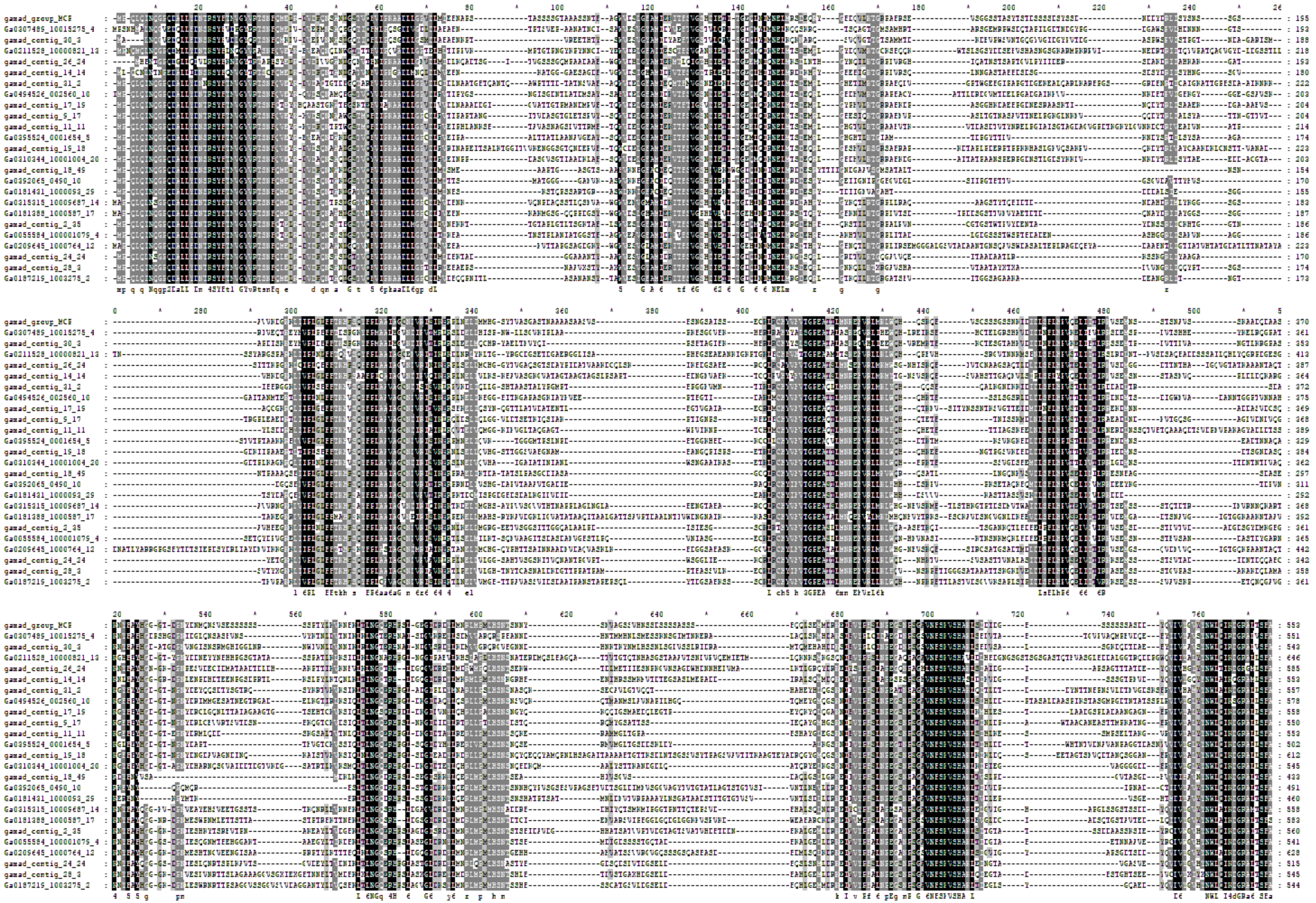

# S3

HHpred search initiated from both MCP alignments brings
mimi-, irido-, asco-, and phycodnavirus MCP with high confidence.

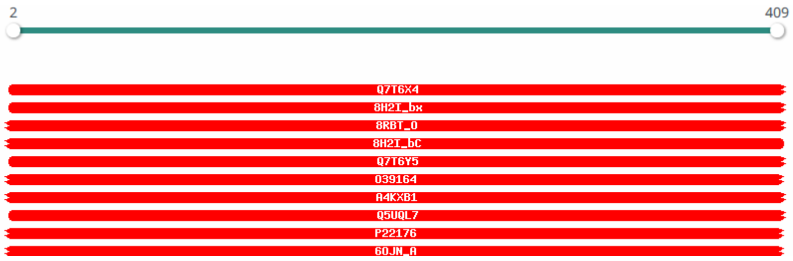


| Nr | Hit | Name | Probability | E-value | Score | SS | Aligned cols | Target Length |
|---|---|---|---|---|---|---|---|---|
| 1 | Q7T6X4 | CAPS2_MIMIV Probable capsid protein 2 OS=Acanthamoeba polyphaga mimivirus OX=212035 GN=MIMI_R439 PE=3 SV=2 | 100 | 2e-74 | 585.41 | 34.7 | 395 | 591 |
| 2 | 8H2I_bx | MCPv1; giant virus, nucleocytoplasmic large DNA viruses (NCLDVs), viral assembly, Paramecium bursaria chlorella virus 1 | 100 | 2.6e-73 | 573.31 | 37 | 397 | 520 |
| 3 | 8RBT_O | Major capsid protein; cryo-EM, subtomogram averaging, EhV-201, enveloped virus, capsid, major capsid protein, VIRUS; 12. | 100 | 1.3e-72 | 565.58 | 39.1 | 390 | 496 |
| 4 | 8H2I_bC | MCPv2; giant virus, nucleocytoplasmic large DNA viruses (NCLDVs), viral assembly, Paramecium bursaria chlorella virus 1 | 100 | 6.9e-73 | 560.36 | 35.9 | 379 | 486 |
| 5 | Q7T6Y5 | CAPS4_MIMIV Probable capsid protein 4 OS=Acanthamoeba polyphaga mimivirus OX=212035 GN=MIMI_R441 PE=3 SV=2 | 100 | 5.7e-73 | 567.88 | 34 | 391 | 512 |
| 6 | O39164 | MCP_IRV16 Major capsid protein OS=Costelytra zealandica iridescent virus OX=68348 GN=MCP PE=3 SV=1 | 100 | 1.1e-72 | 563.02 | 33.3 | 391 | 462 |
| 7 | A4KXB1 | CAPSD_HVAVE Major capsid protein OS=Heliothis virescens ascovirus 3e OX=260797 GN=MCP PE=3 SV=1 | 100 | 9.5e-73 | 561.7 | 32.3 | 391 | 455 |

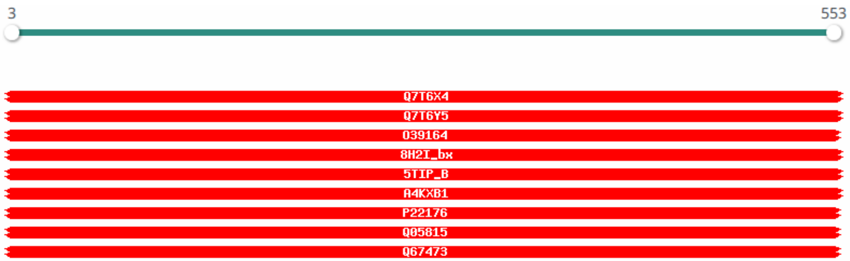


| Nr | Hit | Name | Probability | E-value | Score | SS | Aligned cols | Target Length |
|---|---|---|---|---|---|---|---|---|
| 1 | Q7T6X4 | CAPS2_MIMIV Probable capsid protein 2 OS=Acanthamoeba polyphaga mimivirus OX=212035 GN=MIMI_R439 PE=3 SV=2 | 100 | 1.3e-77 | 669.27 | 40.4 | 419 | 591 |
| 2 | Q7T6Y5 | CAPS4_MIMIV Probable capsid protein 4 OS=Acanthamoeba polyphaga mimivirus OX=212035 GN=MIMI_R441 PE=3 SV=2 | 100 | 1.9e-76 | 651.33 | 44.5 | 425 | 512 |
| 3 | O39164 | MCP_IRV16 Major capsid protein OS=Costelytra zealandica iridescent virus OX=68348 GN=MCP PE=3 SV=1 | 100 | 3e-76 | 643.72 | 44.8 | 434 | 462 |
| 4 | 8H2I_bx | MCPv1; giant virus, nucleocytoplasmic large DNA viruses (NCLDVs), viral assembly, Paramecium bursaria chlorella virus 1 | 100 | 5.4e-76 | 650.01 | 45.8 | 408 | 520 |
| 5 | 5TIP_B | Major capsid protein; VIRAL PROTEIN; HET: 7CV, XYP, FUC, BGC, RM4, MAN, XXR, GLA; 2.0A {Paramecium bursaria Chlorella vi | 100 | 7.3e-76 | 636.15 | 42.6 | 378 | 436 |
| 6 | A4KXB1 | CAPSD_HVAVE Major capsid protein OS=Heliothis virescens ascovirus 3e OX=260797 GN=MCP PE=3 SV=1 | 100 | 1.4e-75 | 637.32 | 44.6 | 411 | 455 |
| 7 | P22176 | MCP_LCDV1 Major capsid protein OS=Lymphocystis disease virus 1 (isolate Darai) OX=654922 GN=MCP PE=3 SV=2 | 100 | 9.2e-76 | 639.42 | 43.3 | 404 | 459 |

*top HHpred hits for Yaraviridae (left) and "Gamadviridae" (right) MCP alignments*

**S4**

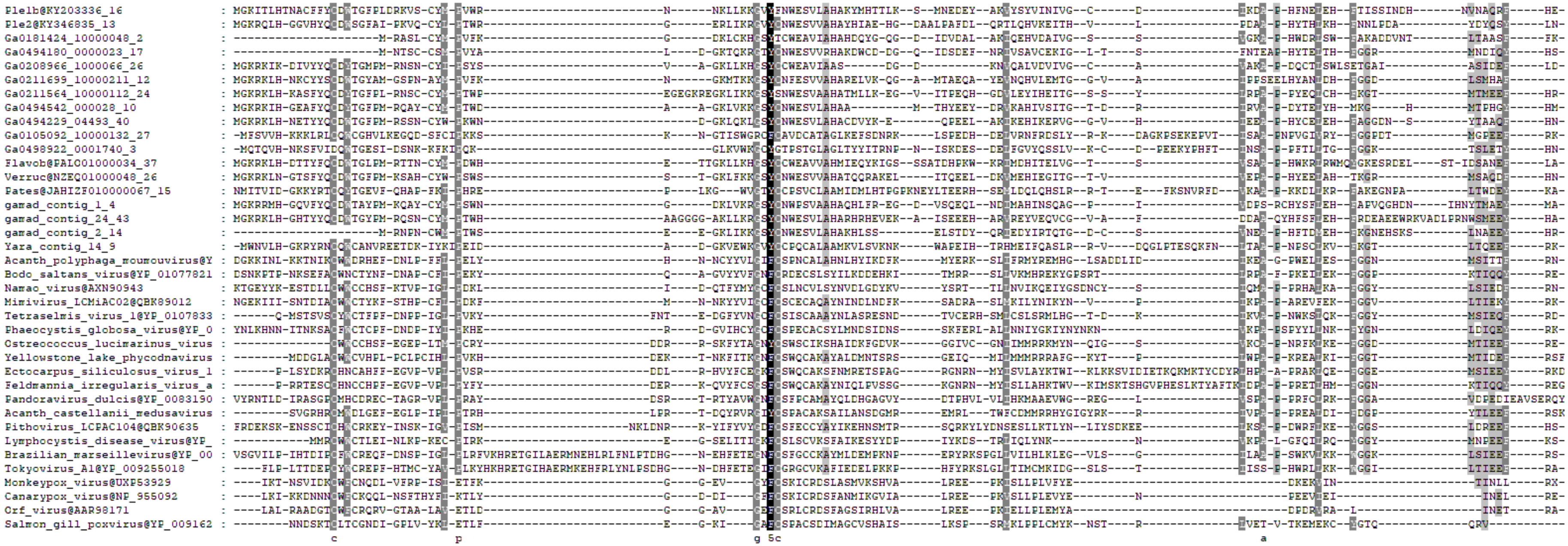

# S5

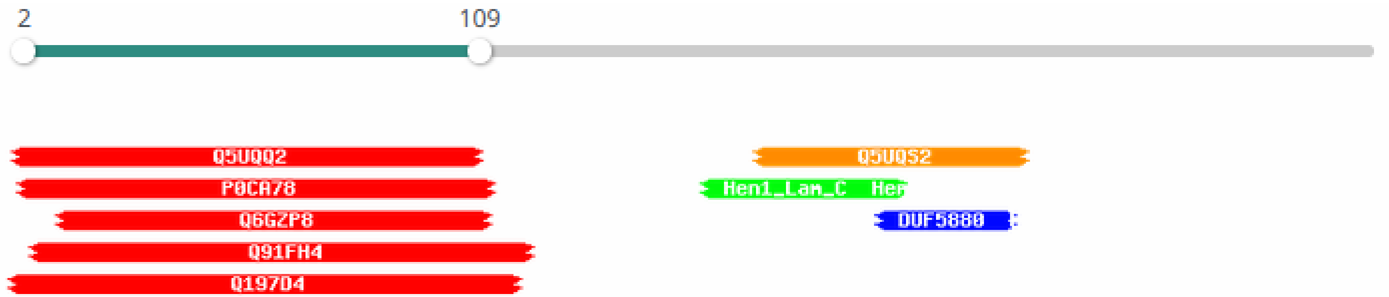


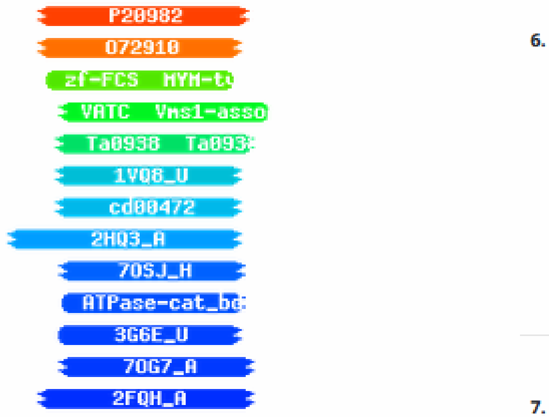


**6. P20982 VLTF2_VACCC Viral late gene transcription factor 2 OS=Vaccinia virus (strain Copenhagen) OX=10249 GN=VLTF2 PE=2 SV=1**

Probability: 94.57%, E-value: 0.1, Score: 47.8, Aligned cols: 44, Identities: 11%, Similarity: 0.121, Template Neff: 3.5

```
Q ss_pred                  CCCCccCCCCCCCCCCCccceeEEEEeCCEEEEEEeeEcCHHHHHHHHHH
Q tmp_VLTF2        8       GKSYYQCDWTGFPMESANCYMPVWSANGKLVWKGSYCNWESVVAHAKE   55 (318)
Q Consensus        8       ~kt~~~CWWC~h~f~~~P~~IP~~~~~g~y~~~G~FCS~eCa~Ayi~~   55 (318)
                           ......||||..+....++.|=+-. +|   ..|.|||--|..++...
T Consensus       46       ~~~~~~CWfC~q~l~~~~~~iETl~-g~---~Vg~FCSkiCrDSfa~~   89 (150)
T P20982         46       NSVIDKCWFCNQDLVFRPISIETYK-GG---EVGYFCSKICRDSLASM   89 (150)
T ss_pred                  CCCCCceecCCCccccCCCeEEEEecC-CC---chheecCHHHHHHHHHH
```

Template alignment

**7. O72910 VLTF2_FOWPN Viral late gene transcription factor 2 OS=Fowlpox virus (strain NVSL) OX=928301 GN=VLTF2 PE=2 SV=1**

Probability: 90.65%, E-value: 0.75, Score: 42.63, Aligned cols: 42, Identities: 14%, Similarity: 0.174, Template Neff: 3.4

```
Q ss_pred                  CCCCccCCCCCCCC-CCCCcceeEEEEeCCEEEEEEeeEcCHHHHHHHH
Q tmp_VLTF2        8       GKSYYQCDWTGFPM-ESANCYMPVWSANGKLVWKGSYCNWESVVAHA   53 (318)
Q Consensus        8       ~kt~~~CWWC~h~f-~~~P~~IP~~~~~g~y~~~G~FCS~eCa~Ayi   53 (318)
                           ......||+|.... ...++.|=+-. +|   ..|.|||-=|..+++
T Consensus       46       ~~~~~~CWfC~q~i~~~~~~~~~iETl~-G~---~IG~FCS~ICrDSfA   88 (154)
T O72910         46       RKDKDNCWFCKQDMNTYNPYFIETLY-GD---HIGVFCSKICRDSFA   88 (154)
T ss_pred                  cCCCCceeccCCcccCCCCeEEEEec-cc---cceeeehHHhHHHHH
```

# S6

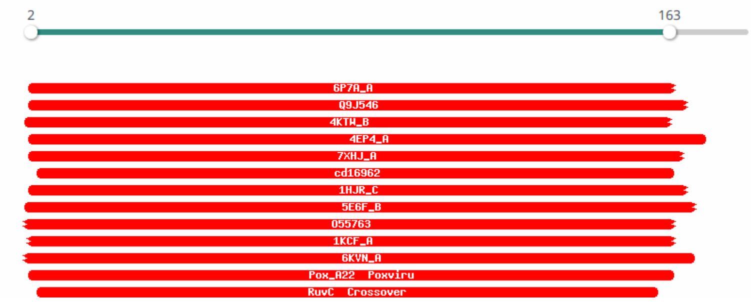


| Nr | Hit | Name | Probability | E-value | Score | SS | Aligned cols | Target Length |
|---|---|---|---|---|---|---|---|---|
| 1 | 6P7A_A | Holliday junction resolvase; RESOLVASE, HYDROLASE; 3.081A {Fowlpox virus} | 99.78 | 4.2e-17 | 115.02 | 14.9 | 138 | 148 |
| 2 | Q9J546 | RUVV_FOWPN Holliday junction resolvase OS=Fowlpox virus (strain NVSL) OX=928301 GN=FPV187 PE=1 SV=1 | 99.78 | 2.2e-17 | 117.98 | 13.3 | 145 | 156 |
| 3 | 4KTW_B | RuvC endonuclease; endonuclease, DNA junctions, replication, recombination, phage packaging, Holliday junction, RNase-H | 99.77 | 5.9e-17 | 116.12 | 15.1 | 141 | 161 |
| 4 | 4EP4_A | Crossover junction endodeoxyribonuclease RuvC; resolvase, Hydrolase; 1.28A {Thermus thermophilus} SCOP: c.55.3.6 | 99.75 | 1.4e-16 | 114.2 | 14.3 | 148 | 166 |
| 5 | 7XHJ_A | Crossover junction endodeoxyribonuclease RuvC; nuclease, DNA repair, Homologous recombination, Holliday junction resolva | 99.74 | 3e-16 | 114.34 | 13.9 | 140 | 179 |
| 6 | cd16962 | RuvC; Crossover junction endodeoxyribonuclease RuvC. Crossover junction endodeoxyribonuclease RuvC is also called Hollid | 99.73 | 4e-16 | 110.27 | 14.1 | 138 | 156 |
| 7 | 1HJR_C | HOLLIDAY JUNCTION RESOLVASE (RUVC); SITE-SPECIFIC RECOMBINASE; 2.5A {Escherichia coli} SCOP: c.55.3.6 | 99.73 | 2.6e-16 | 111.81 | 12.8 | 141 | 158 |
| 8 | 5E6F_B | CNPV261 Holliday junction resolvase protein; Poxvirus, Resolvase, Holliday Junction, Metal-binding, VIRAL PROTEIN; HET: | 99.73 | 1.7e-15 | 107.76 | 16.7 | 146 | 152 |

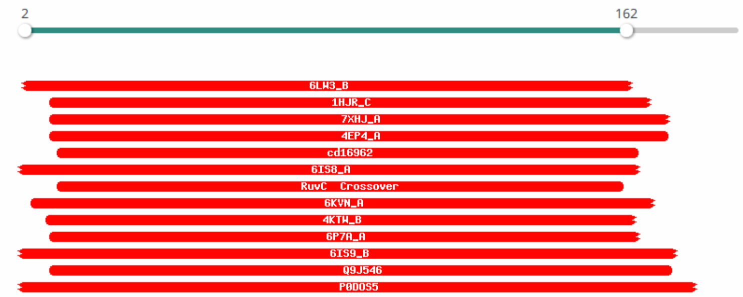


| Nr | Hit | Name | Probability | E-value | Score | SS | Aligned cols | Target Length |
|---|---|---|---|---|---|---|---|---|
| 1 | 6LW3_B | Crossover junction endodeoxyribonuclease RuvC; Holliday junction resolvase RuvC Endonuclease, DNA BINDING PROTEIN; 2.38A | 99.68 | 1.4e-14 | 101.5 | 16 | 147 | 163 |
| 2 | 1HJR_C | HOLLIDAY JUNCTION RESOLVASE (RUVC); SITE-SPECIFIC RECOMBINASE; 2.5A {Escherichia coli} SCOP: c.55.3.6 | 99.68 | 1.4e-14 | 101.01 | 15.7 | 145 | 158 |
| 3 | 7XHJ_A | Crossover junction endodeoxyribonuclease RuvC; nuclease, DNA repair, Homologous recombination, Holliday junction resolva | 99.67 | 1.9e-14 | 102.89 | 16 | 151 | 179 |
| 4 | 4EP4_A | Crossover junction endodeoxyribonuclease RuvC; resolvase, Hydrolase; 1.28A {Thermus thermophilus} SCOP: c.55.3.6 | 99.65 | 7.1e-14 | 98.46 | 16.8 | 152 | 166 |
| 5 | cd16962 | RuvC; Crossover junction endodeoxyribonuclease RuvC. Crossover junction endodeoxyribonuclease RuvC is also called Hollid | 99.65 | 5e-14 | 97.77 | 15.4 | 142 | 156 |
| 6 | 6IS8_A | Monokaryotic chloroplast 1; Holliday junction resolvase-DNA complex, DNA BINDING PROTEIN-DNA complex; HET: 1PE; 1.68A {Z | 99.61 | 1.1e-13 | 98.5 | 14.5 | 151 | 174 |
| 7 | PF02075.21 | ; RuvC ; Crossover junction endodeoxyribonuclease RuvC | 99.6 | 1.8e-13 | 94.57 | 14.4 | 137 | 148 |
| 8 | 6KVN_A | NtMOC1; chloroplast, resolvase, PLANT PROTEIN; 2.02A {Nicotiana tabacum} | 99.6 | 2.7e-13 | 96.23 | 15.3 | 150 | 171 |

*top HHpred hits for “Gamadviridae” (right) and Yaraviridae (left) RuvC alignments*

# S7

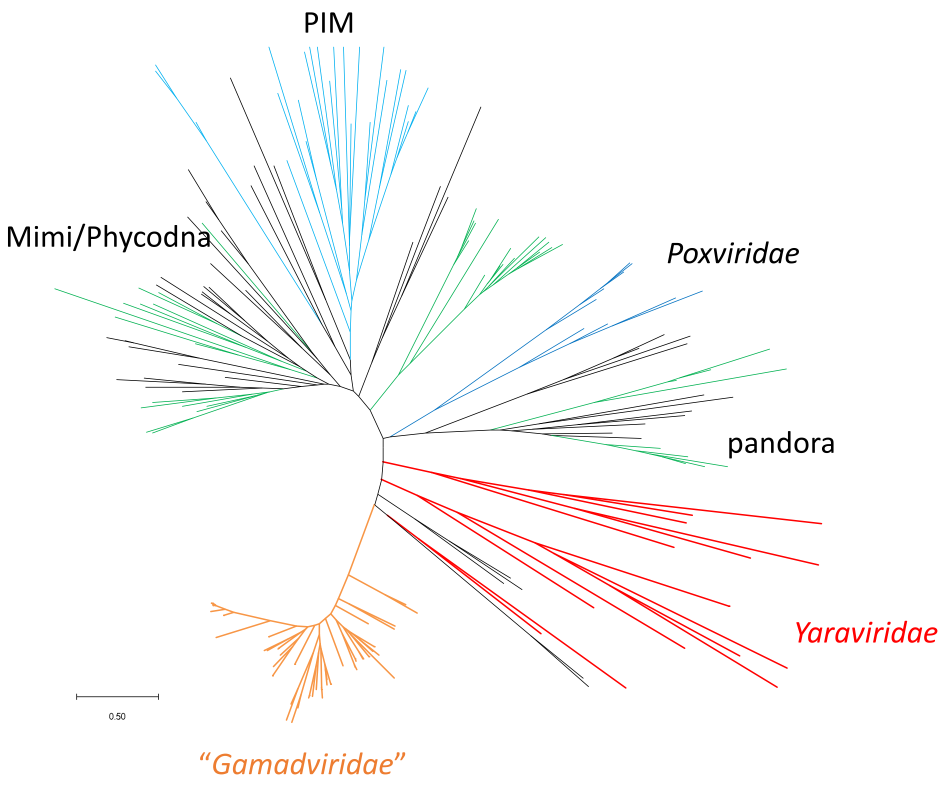

PIM
Mimi/Phycodna
*Poxviridae*
pandora
*Yaraviridae*
0.50
*"Gamadviridae"*

# S8

#### SF3_hel1 (most of *Yaraviridae*; Yaravirus gene 69)

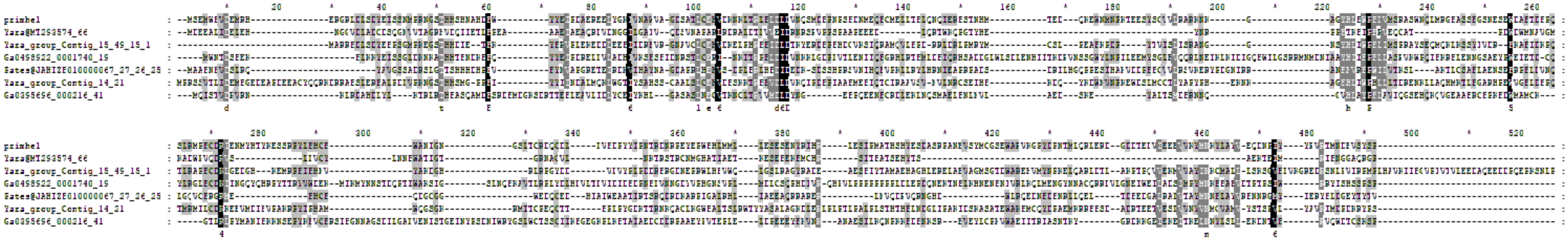

##### primase domain

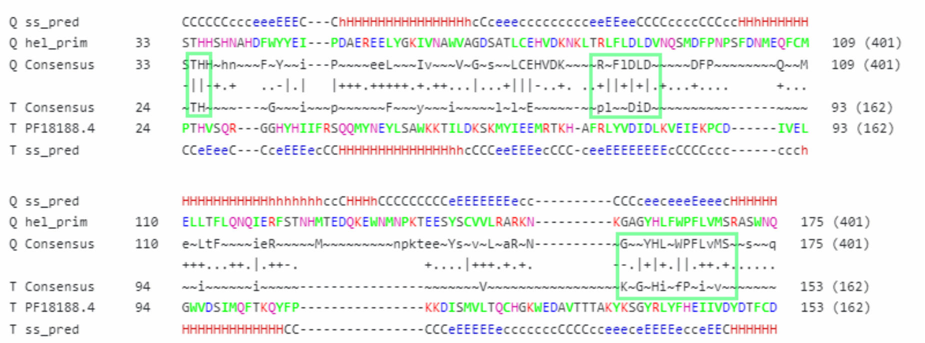

##### helicase domain

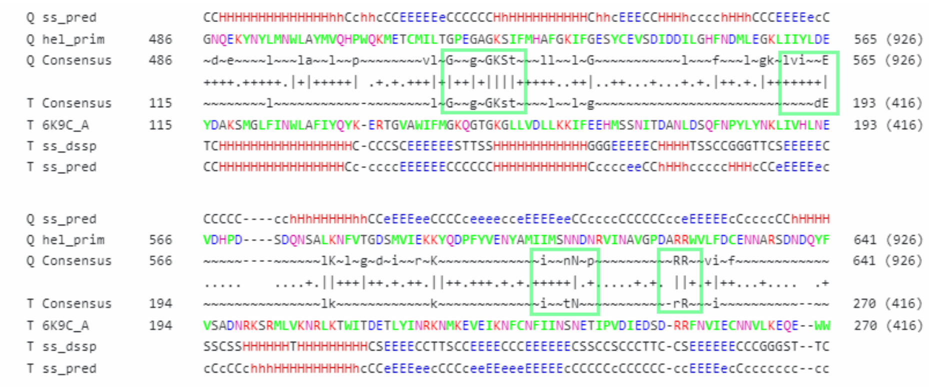

### SF3_hel1 (Yaravirus gene 69)

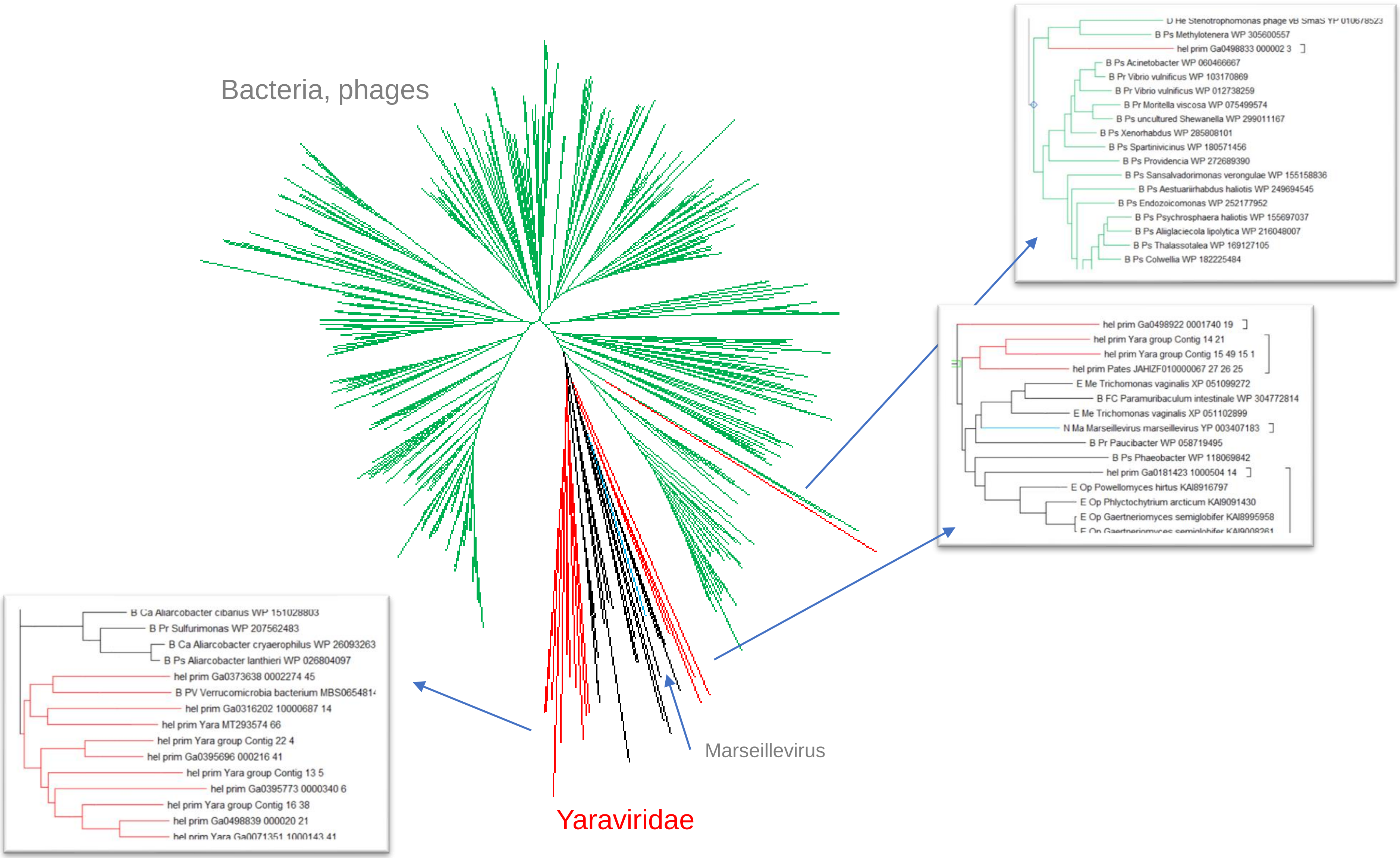

# S10

SF3_hel2

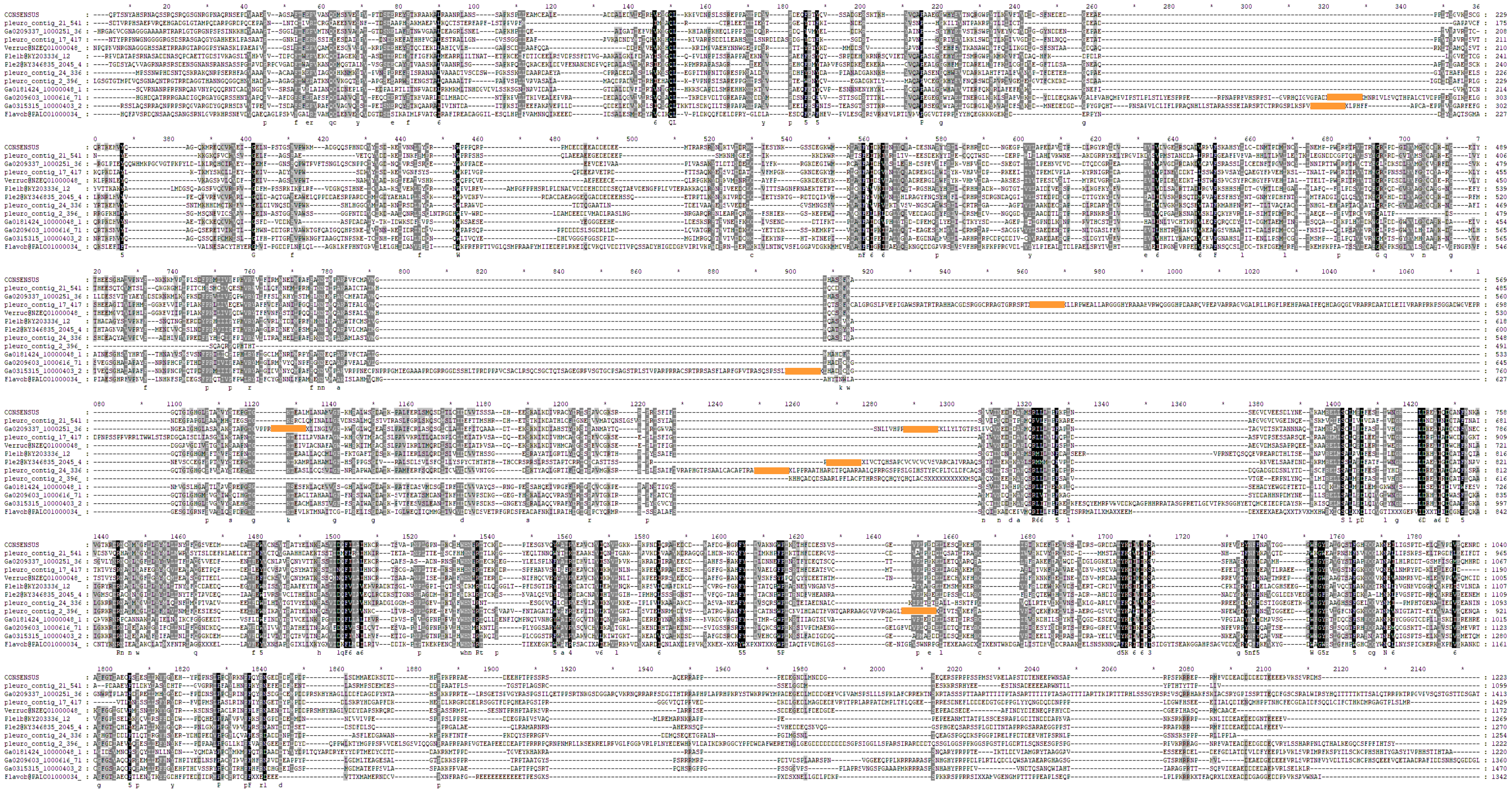

some SF3_hel2 proteins have frameshifts (marked in orange on the figure).

# S11

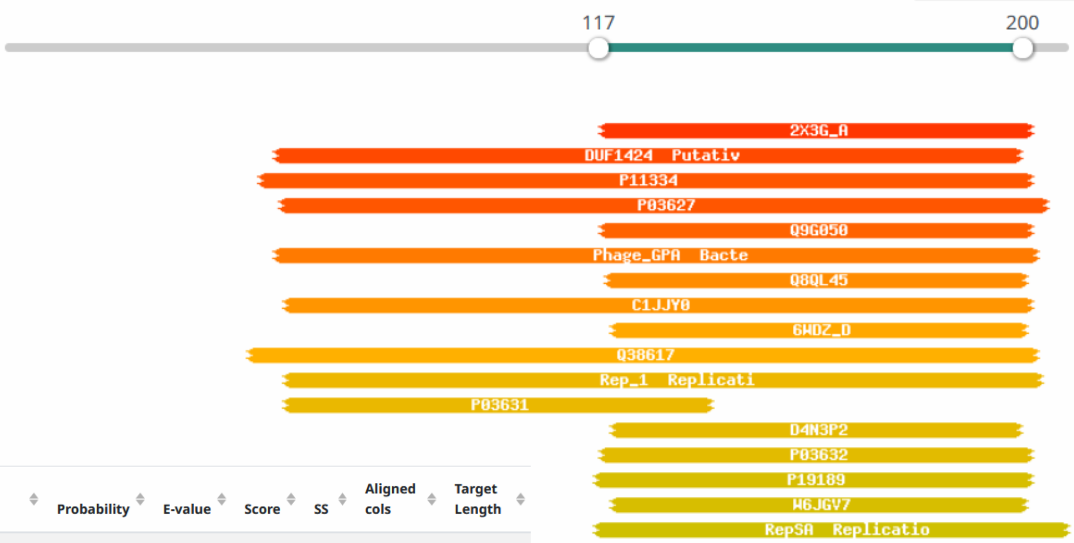


| Nr | Hit | Name | Probability | E-value | Score | SS | Aligned cols | Target Length |
|---|---|---|---|---|---|---|---|---|
| 1 | 2X3G_A | SIRV1 HYPOTHETICAL PROTEIN ORF119; VIRAL PROTEIN, HOST MEMBRANE; HET: MSE; 1.8A {SULFOLOBUS ISLANDICUS RUDIVIRUS 1 VARIA | 95.52 | 0.097 | 42.87 | 6 | 65 | 119 |
| 2 | PF07232.14 | ; DUF1424 ; Putative rep protein (DUF1424) | 93.77 | 6.8 | 36.39 | 13.8 | 115 | 316 |
| 3 | P11334 | REP_SPV4 Replication-associated protein ORF2 OS=Spiroplasma virus 4 OX=10855 GN=ORF2 PE=3 SV=1 | 92.74 | 3.1 | 39.96 | 10.4 | 107 | 320 |
| 4 | P03627 | REP_BPPF3 Replication-associated protein G2P OS=Pseudomonas phage Pf3 OX=10872 GN=II PE=3 SV=1 | 92.66 | 8.1 | 36.1 | 12.6 | 109 | 278 |
| 5 | Q9G050 | REP_BPPHM Replication-associated protein VP4 OS=Bdellovibrio phage phiMH2K OX=145579 GN=ORF4 PE=3 SV=1 | 91.61 | 1.4 | 42.09 | 6.8 | 67 | 315 |
| 6 | PF05840.16 | ; Phage_GPA ; Bacteriophage replication gene A protein (GPA) | 89.69 | 12 | 35.26 | 10.7 | 119 | 316 |
| 7 | Q8QL45 | Y117_SIRV1 Uncharacterized protein 117 OS=Sulfolobus islandicus rod-shaped virus 1 OX=157898 GN=117 PE=4 SV=1 | 88.17 | 2.8 | 35.65 | 5.1 | 63 | 117 |

S12

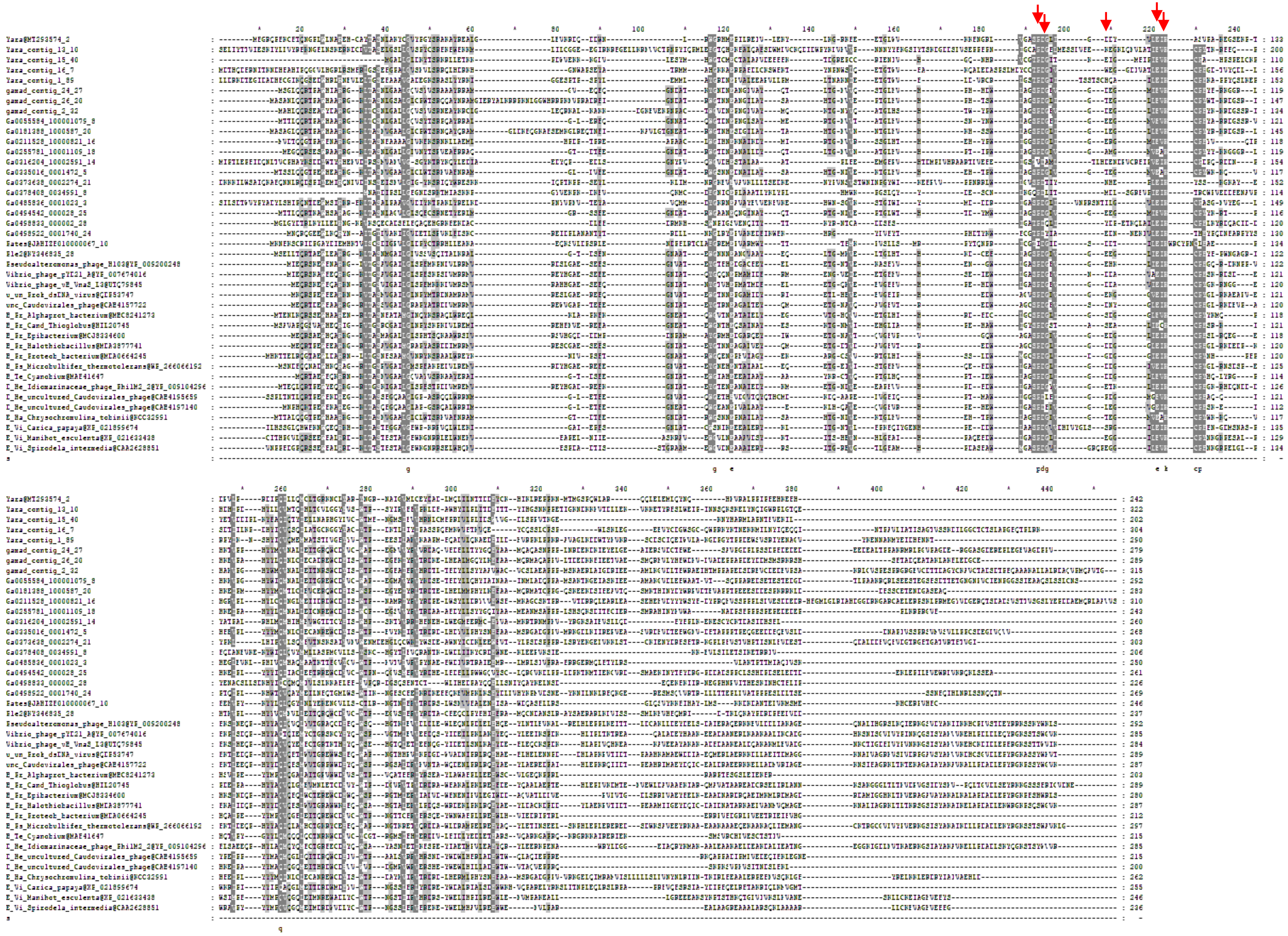

# S13

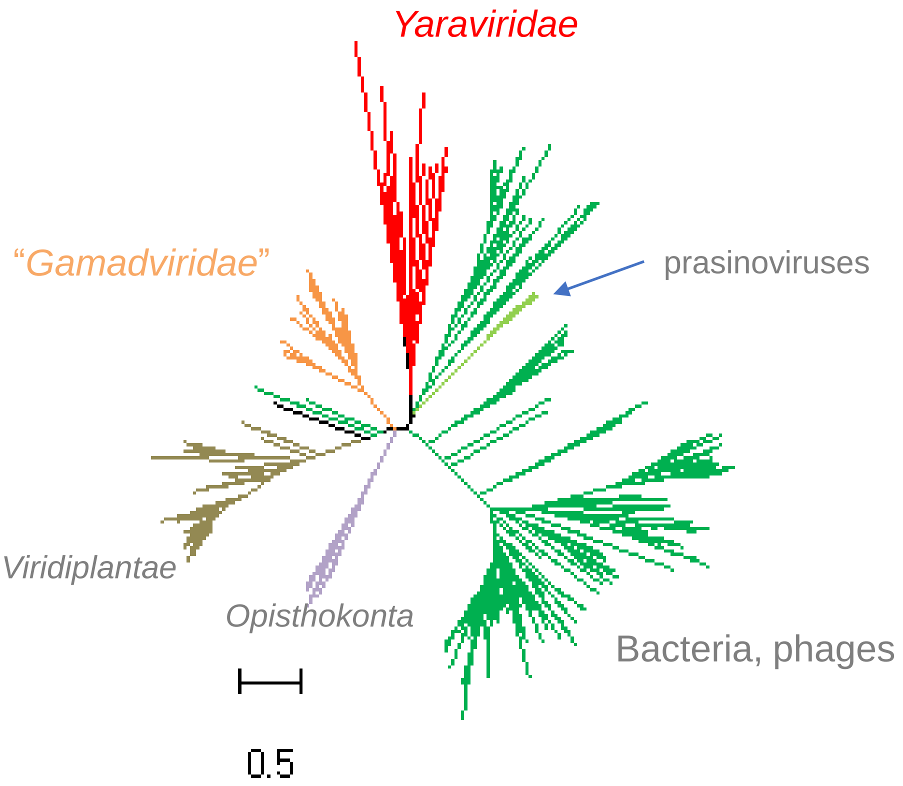
Yaraviridae
"Gamadviridae"
prasinoviruses
Viridiplantae
Opisthokonta
Bacteria, phages
0.5

# S14

#### gene48 with nr hits

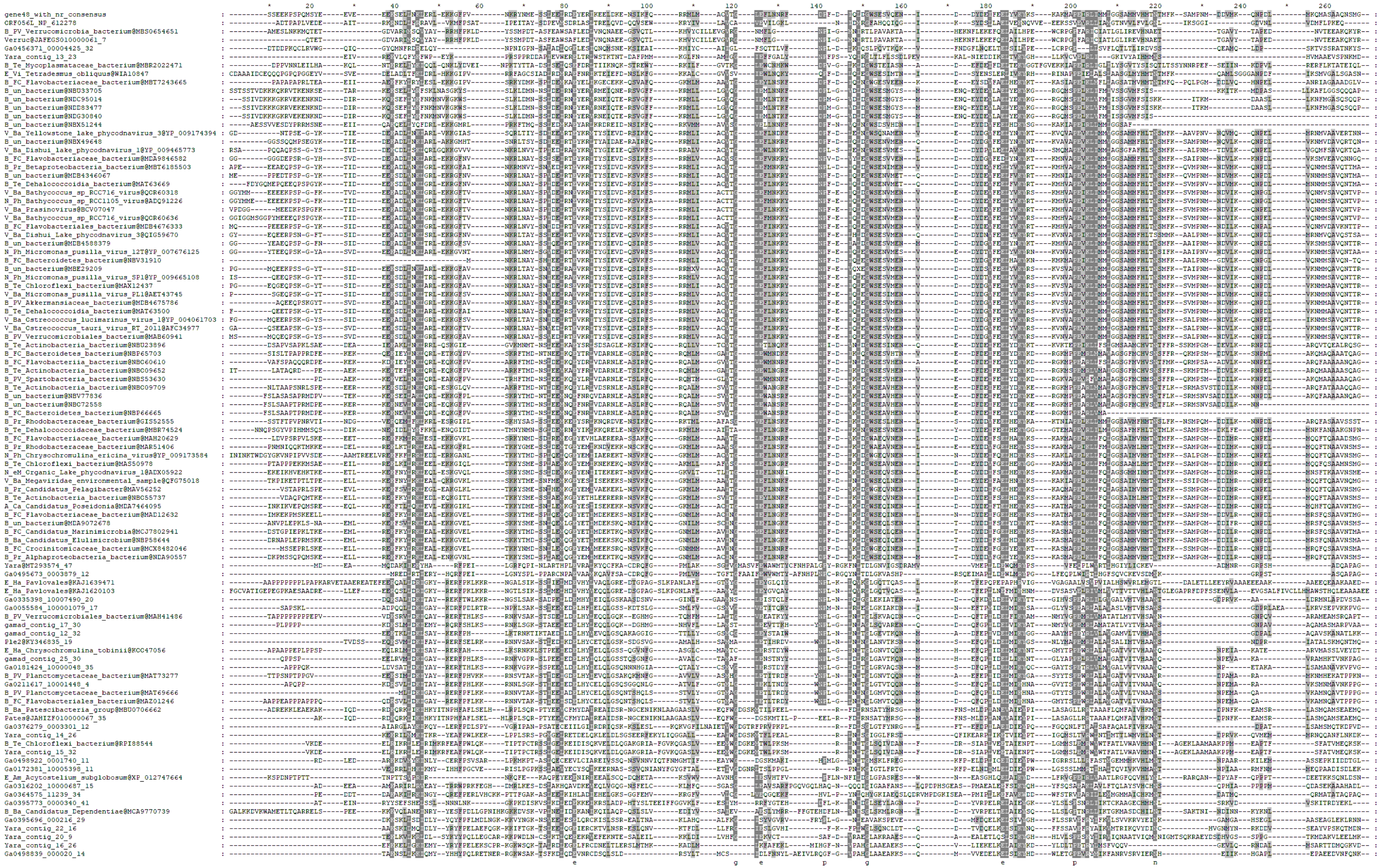

A

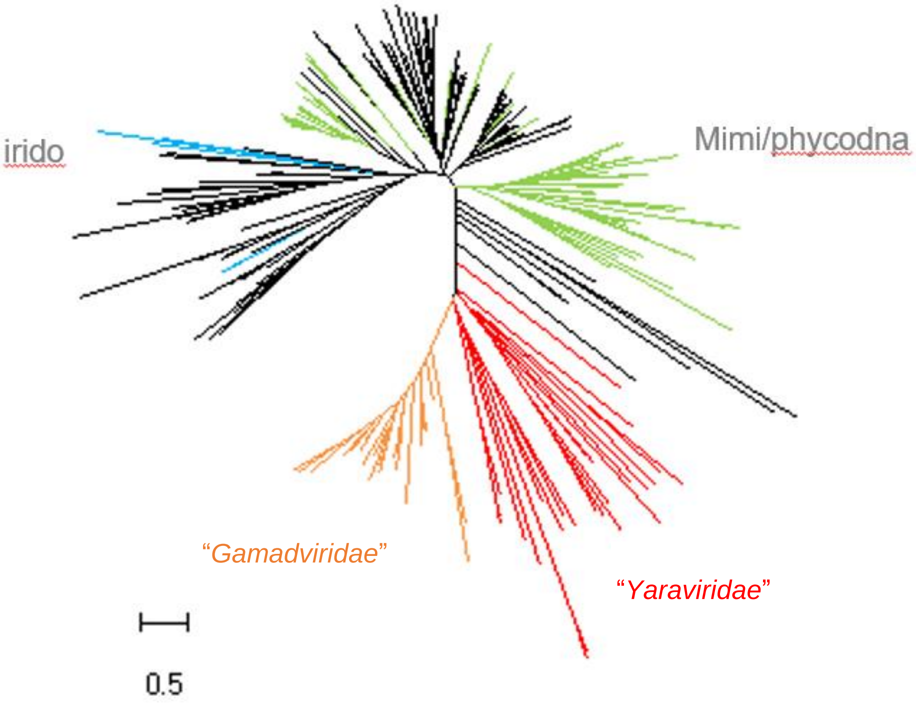


B

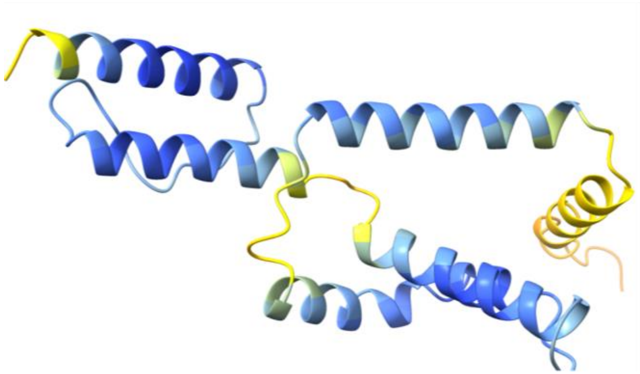

gene48 (Ple2@KY346835_19)

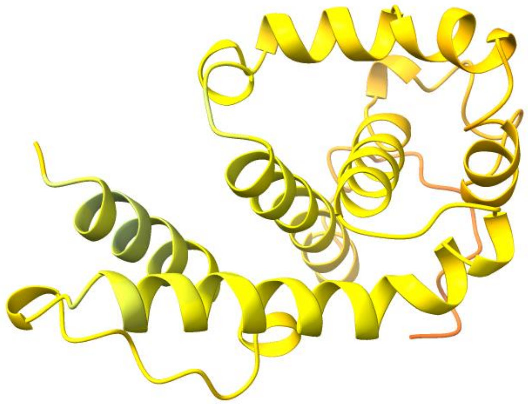

(Yaravirus)

# S16

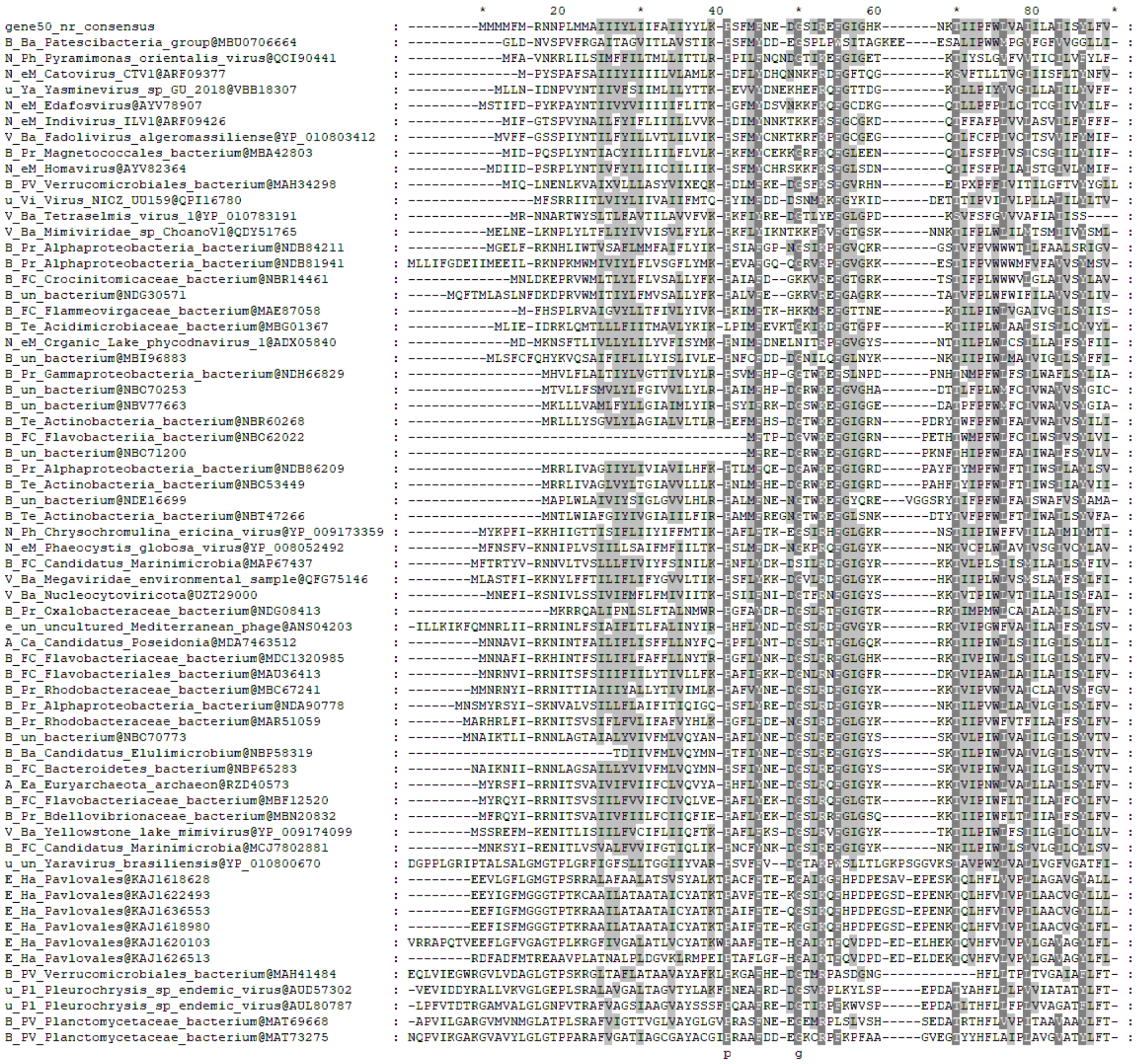

Mriya_50 with nr hits

# S17

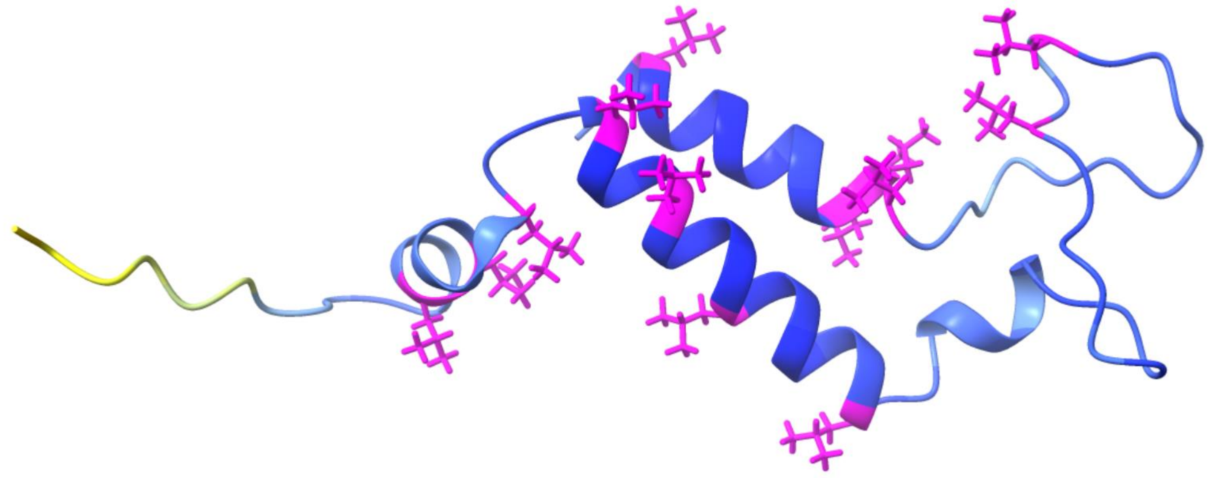

Mriya_50 (Ple2@KY346835_21)

# S18

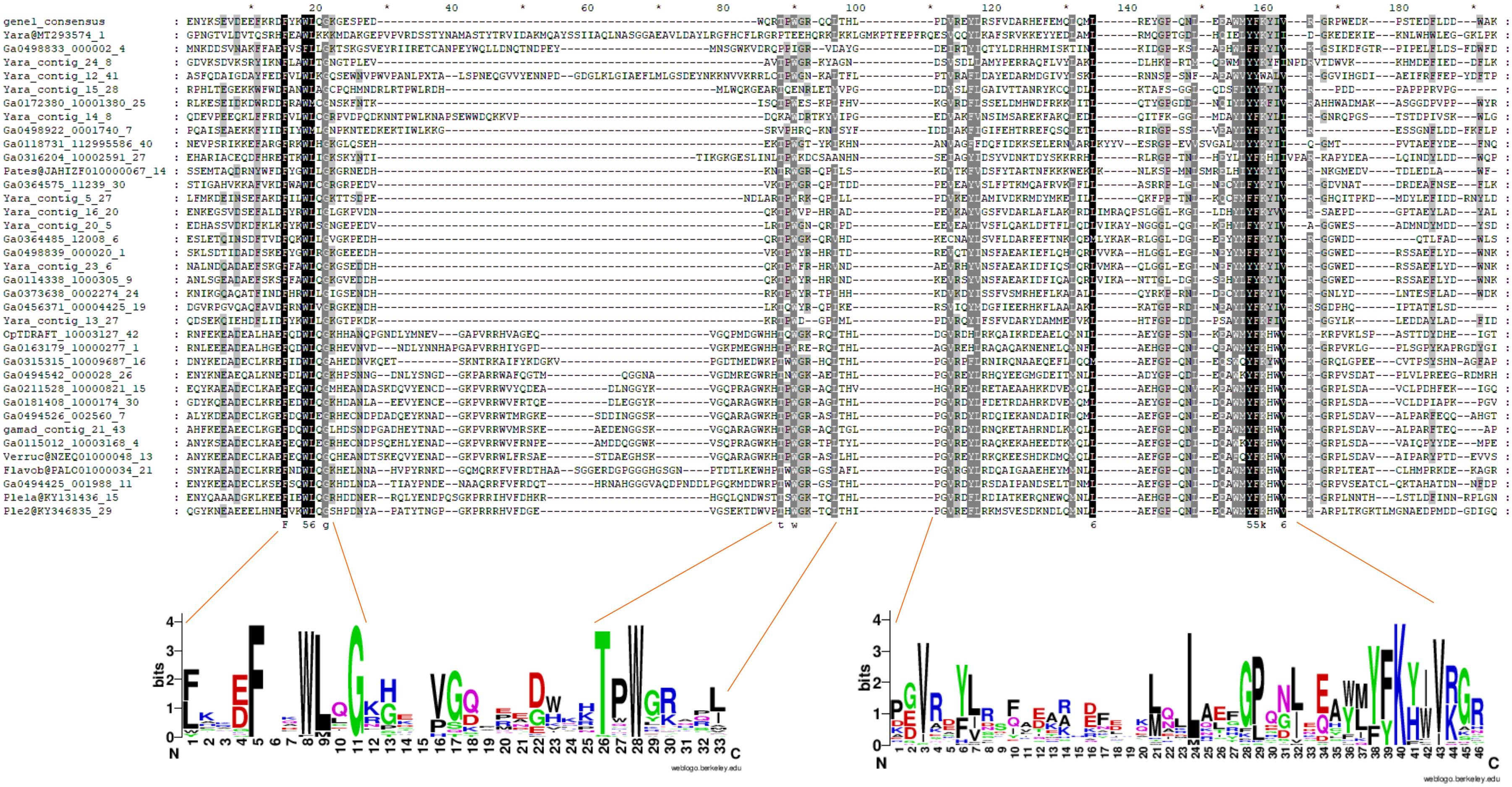

# S19

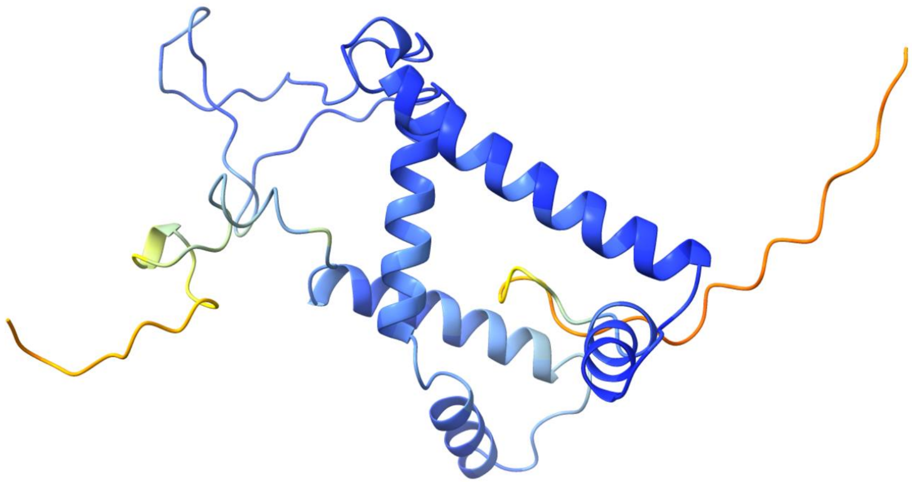

Mriya_1, exemplified with Ple2@KY346835_29

Mriya_51

**S20**

A

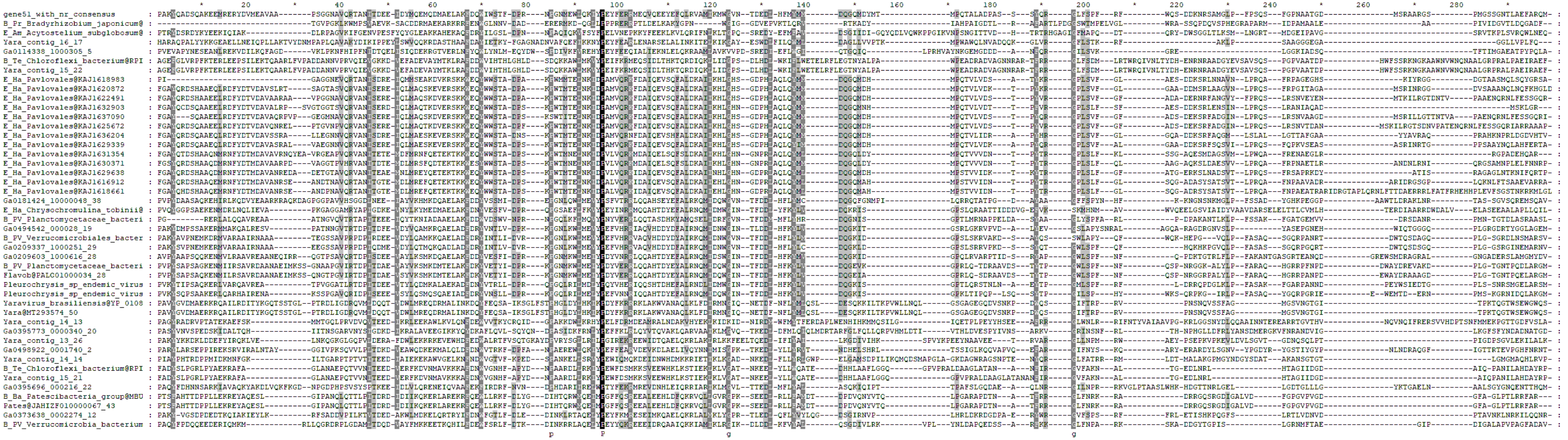

B

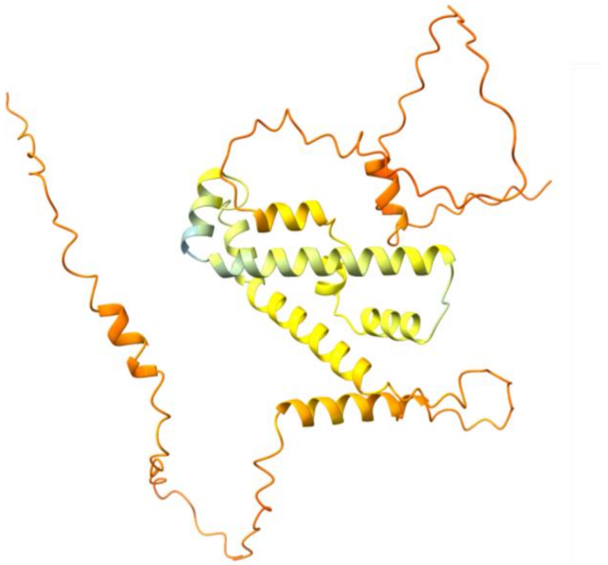

(Yara@MT293574_50, aka YP_010800671)

**S21**

A

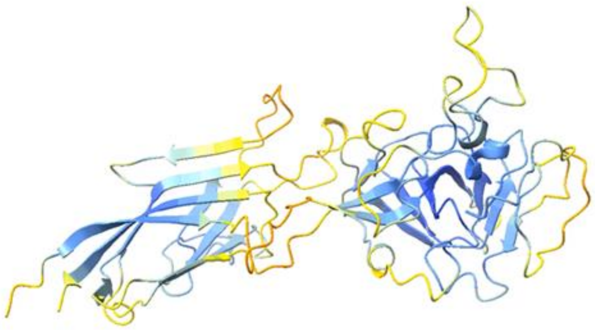

Foldseek:
4 uncharacterized proteins

Dali
6.4 , 6g41-A , MOLECULE: MINOR CAPSID PROTEIN;
→ Cafeteriavirus-dependent mavirus (Varidnaviria, Bamfordvirae, Preplasmiviricota)
5.4 , 3j26-N , MOLECULE: CAPSID PROTEIN V20;
→ resolution structure of the Sputnik virophage by cryo-EM

B

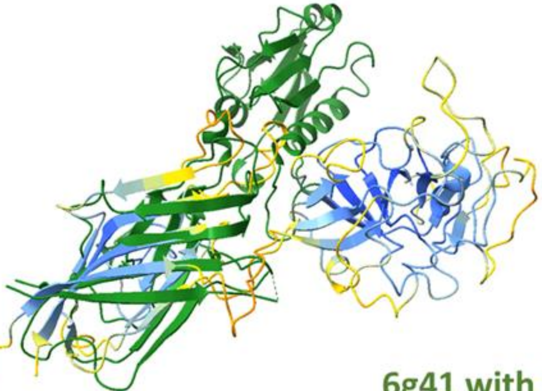

6g41 with SJR (overlayed with Ple_22) and insertion domain

# S22

PEV_26 - ssb

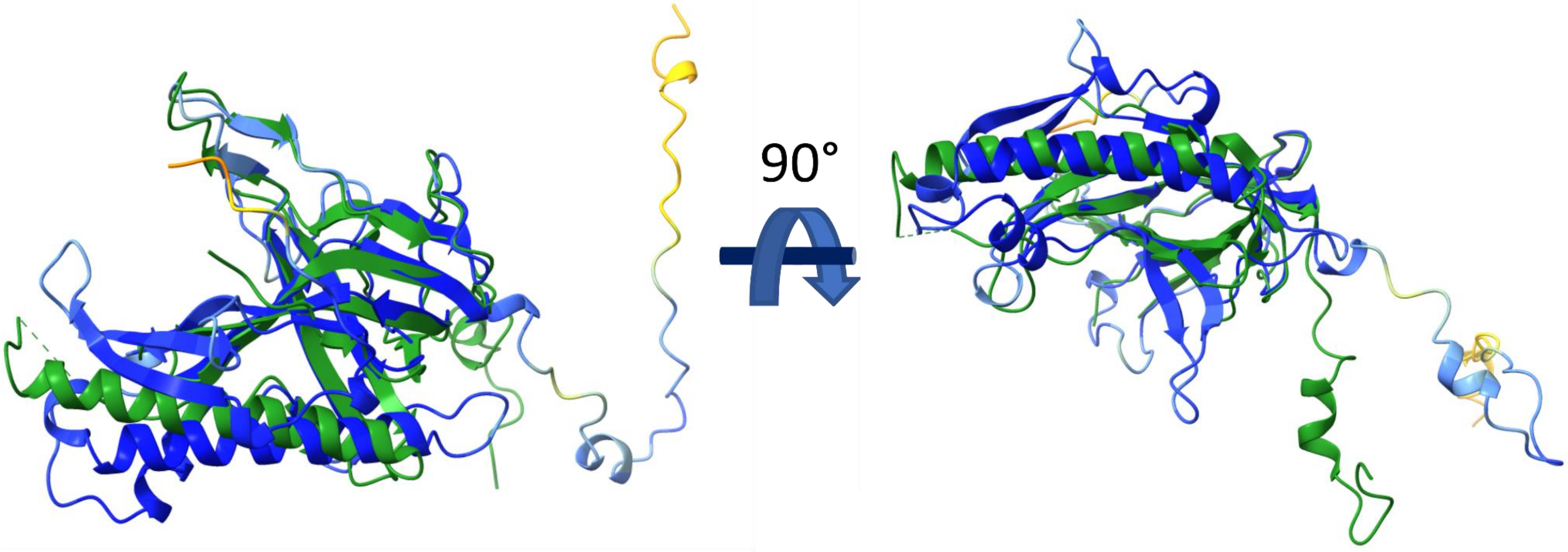

# S23

Yaravirus  minor capsid protein (YP_010800666.1)

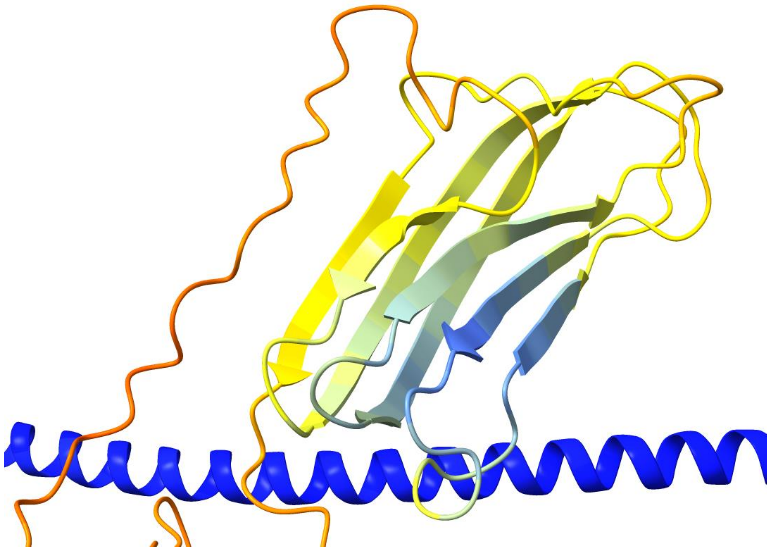

# S24

Yaravirus minor capsid protein (C-terminal of YP_010800666.1)

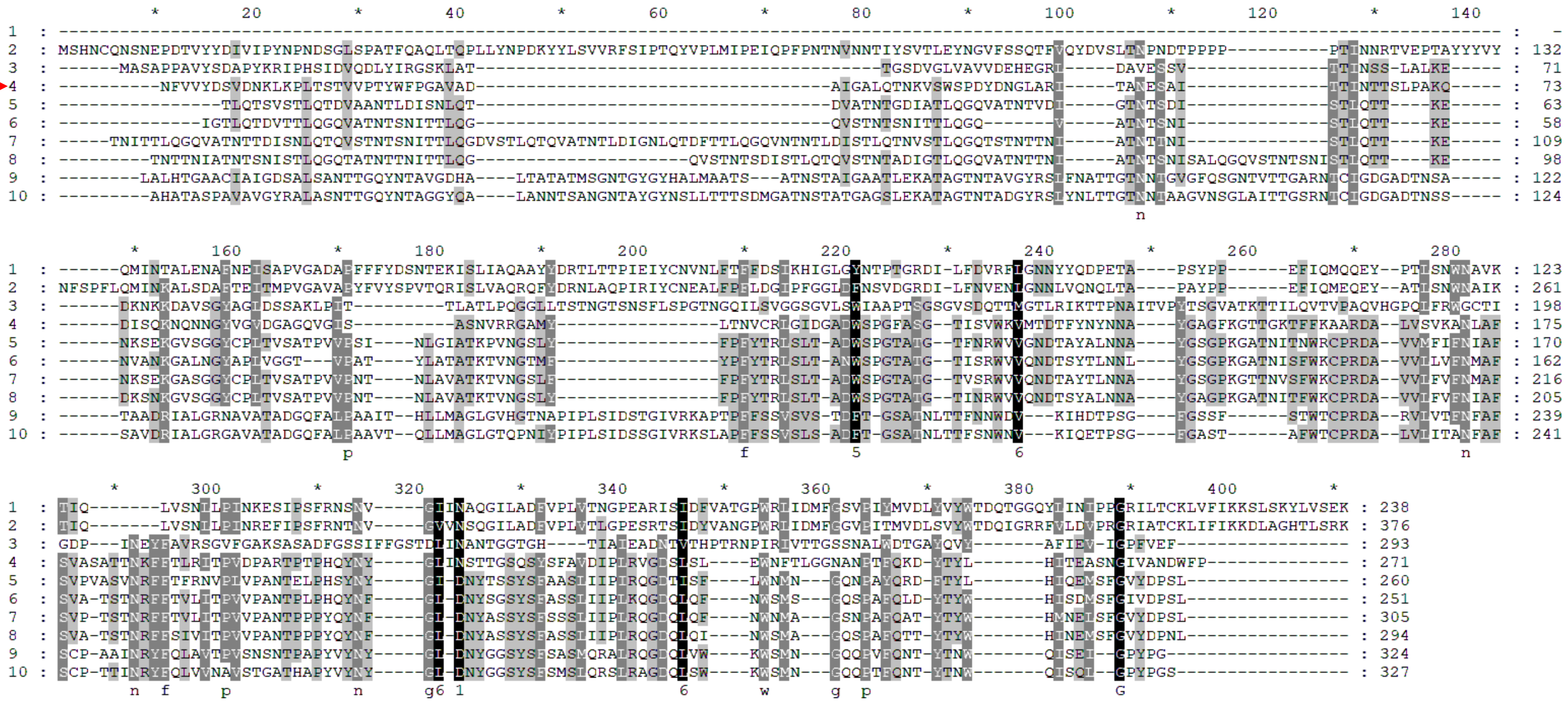

1 Sputnik_UMZ08530 .1 minor virion protein [Mimivirus-dependent virus Sputnik]
2 Zamilon_YP_008859634 .1 minor virion protein [Zamilon virus]
3 YP_010800667.1 hypothetical protein QKS73_gp57 [Yaravirus brasiliensis]
4 YP_010800666.1:200-470 tail needle DNA stabilization protein [Yaravirus brasiliensis]
5 YP_009507121.1:265-524 chromosome segregation protein (SMC) [Tunisvirus fontaine2]
6 YP_009094634.1:305-555 hypothetical protein MEL_133 [Melbournevirus]
7 YP_009345180.1:241-545 hypothetical protein B1750_gp035 [Noumeavirus]
8 YP_009238991.1:321-614 hypothetical protein A3303_gp486 [Brazilian marseillevirus]
9 YP_010804828.1:961-1284 hypothetical protein QKU73_gp304 [Acanthamoeba castellanii medusavirus]
10 YP_010802084.1:451-777 hypothetical protein QKT49_gp315 [Acanthamoeba castellanii medusavirus]
